# Large-scale Phenomic and Genomic Analysis of Brain Asymmetrical Skew

**DOI:** 10.1101/756395

**Authors:** Xiang-Zhen Kong, Merel Postema, Amaia Carrión Castillo, Antonietta Pepe, Fabrice Crivello, Marc Joliot, Bernard Mazoyer, Simon E. Fisher, Clyde Francks

**Author notes:** **Correspondence Author:** Clyde Francks, D.Phil., Wundtlaan 1, 6525 XD Nijmegen, The Netherlands.

## Abstract

The human cerebral hemispheres show a left-right asymmetrical torque pattern, which has been claimed to be absent in chimpanzees. The functional significance and developmental mechanisms are unknown. Here we carried out the largest-ever analysis of global brain shape asymmetry in magnetic resonance imaging data. Three population datasets were used, the UK Biobank (*N* = 39,678), Human Connectome Project (*N* = 1,113) and BIL&GIN (*N* = 453). At the population level, there was an anterior and dorsal skew of the right hemisphere, relative to the left. Both skews were associated independently with handedness, and various regional grey and white matter metrics oppositely in the two hemispheres, as well as other variables related to cognitive functions, sociodemographic factors, and physical and mental health. The two skews showed SNP-based heritabilities of 4-13%, but also substantial polygenicity in causal mixture model analysis, and no individually significant loci were found in GWAS for either skew. There was evidence for a significant genetic correlation (*r*_*g*_=−0.40, *p*=0.0075) between horizontal brain skew and Autism Spectrum Disorder. These results provide the first large-scale description of population-average brain skews and their inter-individual variations, their replicable associations with handedness, and insights into biological and other factors which associate with human brain asymmetry.

## Introduction

A counter-clockwise twist of the whole brain along the anterior-posterior axis, i.e. the fronto-occipital torque, has been widely reported in humans since observations in the middle of the 20th century (e.g., [1–6]; see [7] for a review). This global twisting is manifested by several features, including a more anteriorly protruding right frontal lobe (frontal petalia) and posteriorly protruding left occipital lobe (occipital petalia), a so-called “bending” of the right frontal and left occipital lobes across the midline, and relative increases in the dimensions (e.g., volume and width) of the right frontal and left occipital poles [7].

Torque has recently been reported to be mostly or wholly absent in our closest living relatives the chimpanzees [8–10], although some studies have reported torque in chimpanzees [11] and other primates [10, 12, 13]. There is also evidence for alterations of torque in cognitive and neuropsychiatric disorders, including developmental stuttering [14], dyslexia [15], schizophrenia ([16–18]; but see [19]), attention-deficit/hyperactivity disorder [20], and depression [21, 22]. While the sample sizes were not large in these previous studies (e.g., 37 cases and 44 controls in [18]; 231 cases and 68 controls in [21]), and further replication is needed, these results suggest that the global brain asymmetry pattern may reflect an optimal organization of the human brain, and deviation from it might serve as a biomarker of brain dysfunction.

Besides torque on the anterior-occipital axis, asymmetry on the dorsal-ventral axis has also been reported, but less consistently or well described. An early study reported that the left hemisphere was shifted dorsally relative to the right [23], but recent work based on MRI analysis of 91 human brains found the opposite pattern, i.e. the left hemisphere shifted significantly downward relative to the right [8]. Again, this pattern was reported to be human specific, in comparison to chimpanzees [8]. Variation in ‘vertical’ asymmetry has not been linked to behavioural differences, or disorder risk, as far as we are aware. In addition, neither vertical nor horizontal asymmetries have been measured in large-scale population analysis in thousands of people, to assess their averages, variances or correlations.

It was posited in 1874 that “difference of [brain] structure of necessity implies difference in function” [24]. However, it has proven surprisingly difficult to link brain structural asymmetries to lateralized functions [25–28]. For example, handedness is one of the most clearly evident functional lateralizations, such that in the general population roughly 90% of people are right-handed, and 10% left-handed [29, 30]. In a series of studies based on X-Ray Computed Tomography (CT), LeMay and colleagues reported that the right occipital lobe was more often wider than the left in left-handers, which was the opposite of that found in right-handers [2, 31–33]. Some researchers have even attempted to use the asymmetrical anatomy of skull endocasts to infer handedness in hominins [2, 34, 35].

However, other investigations of the relationships between handedness and features of global brain asymmetry, including more recent studies with up-to-date methodology, have produced inconsistent or negative findings [36–39]. It has therefore been noted that handedness and global brain asymmetry might not be associated at all, and in any case, their relationship is clearly far from absolute [37, 38, 40, 41]. Several MRI studies of regional structural asymmetries that may partly reflect global asymmetry, such as cortical thickness asymmetry of frontal and occipital regions, have also failed to find associations with handedness [5, 6, 42, 43]. Overall, the mixed results may reflect differences in many factors, including limited imaging quality in early studies, statistical power related to small sample sizes, and potential biases when measuring global asymmetry, perhaps especially for manual approaches. A large-scale survey using high-resolution imaging, and objective analysis, is therefore needed to understand the relevance of global brain asymmetry to handedness.

Population-level, average left-right differences of global brain anatomy suggest a genetic-developmental program that is inherently lateralized [44–47]. Torque has been observed in fetal brains by the second trimester of pregnancy [4], as have other region-specific brain asymmetries [44], which further supports a genetic influence. McShane et al. reported left-right differences of occipital petalia and width that were related to ethnic origin, suggesting genetic contributions to variability [48]. A recent, large-scale population study indicated a torque-like pattern of cortical thickness asymmetry, with frontal regions being generally thicker on the left hemisphere, and occipital regions thicker on the right [6]. In the same study, twin and/or family-based analysis found heritabilities of up to roughly 20% for some of these regional cortical thickness asymmetry measures, for example in the lateral occipital and rostral middle frontal regions, which again suggests that genetic variability may affect global brain asymmetries. A study of vervet monkeys [12] also reported heritabilities of 10%-30% for measures of global brain asymmetry, which were methodologically very similar to those used in the present study, i.e. based on skewing brain MRI data in order to register to a symmetrical template (see below).

Recent large-scale genome-wide association studies (GWAS) have identified specific loci involved in regional brain asymmetries [49–51], and also in left-handedness [52–54]. Microtubule-related genes have been particularly implicated by these studies, which is consistent with a role of the cytoskeleton in setting up cellular chirality during embryonic development of the left-right axis of other organs in other species [55–58]. However, the specific genes involved in global brain asymmetry remain unknown. Early life factors that are known to influence handedness such as birthweight and multiple birth [29, 59] could also contribute to global brain asymmetry, but this has not previously been studied in large population data.

Here, we present the largest-ever analysis of global brain asymmetry, in three independent datasets: the UK Biobank (*N* = 39,678), Human Connectome Project (HCP, *N* = 1,113) and BIL&GIN (*N* = 453, roughly balanced for left/right handedness). First, the two components of global brain asymmetry, i.e. the horizontal and vertical asymmetry skews, were extracted from brain MRI data for each individual, to capture global left-right differences along the anterior-posterior and dorsal-ventral axes. The population distributions of these two measures were examined in the three datasets, to clarify the population-level average direction and variance of each asymmetry component. In addition, the reliability of these two measures was confirmed using a test-retest dataset from the HCP (30 subjects in the dataset had two scans each). Next, we investigated the relationships of horizontal and vertical skews with handedness in each of the datasets. Then, we used extensive phenotypic data in the UK Biobank, including early life factors, sociodemographic factors, regional grey and white matter measures derived from brain MRI, and variables related to cognitive functions and health, to explore other potential correlates of global brain asymmetry. Furthermore, we estimated the heritabilities of the horizontal and vertical skews using genome-wide genotype data in the UK Biobank data, and twin data from the HCP, and also investigated their genetic correlations with other traits and disorders. We also used the UK Biobank data to screen the genome for single nucleotide polymorphisms (SNPs) that associate with the horizontal or vertical brain skew measures, at the single-SNP, gene and pathway levels.

## Materials and Methods

### Datasets

#### UK Biobank

Data were obtained from the UK Biobank as part of research application 16066, with Clyde Francks as the principal applicant. This is a general adult population cohort. The data collection in the UK Biobank, including the consent procedure, has been described elsewhere [60]. Informed consent was obtained by the UK Biobank for all participants. For this study, we used data from the February 2020 release of 39,678 participants’ brain T1-wighted MRI data, after bias field correction and brain extraction (i.e. *T1_unbiased_brain.nii.gz*) [61]. The median age of the 39,678 subjects was 64 years, range 44 to 82 years, and 20,998 subjects were female. Handedness was assessed based on responses to the question: “*Are you right or left handed?*” with four response options: “*Right-handed*”, “*Left-handed*”, “*Use both right and left equally*”, and “*Prefer not to answer*”. Those who preferred not to answer were excluded for association analysis of global brain asymmetry with handedness, leaving 35,338 right-handers, 3,712 left-handers, and 614 ‘ambidextrous’ with brain asymmetry measures. We also made use of genome-wide genotype data for single nucleotide polymorphisms (SNPs) as described previously [62], as well as other phenotypic data, including early life factors, sociodemographic factors, regional grey and white matter measures that had already been derived from the brain imaging images, and variables related to cognitive functions and health. Information on these additional phenotype measures are available via the Data Showcase on the UK Biobank website (https://www.ukbiobank.ac.uk/). Our study made use of imaging-derived phenotypes generated by an image-processing pipeline developed and run on behalf of UK Biobank [61].

#### Human Connectome Project

The HCP comprises 1,113 individuals with MRI data (606 females, age range 22-37 years at the time of scanning) of varying ethnicities (https://humanconnectome.org/). The HCP contains 143 monozygotic twin pairs and 85 dizygotic twin pairs, as well as unrelated individuals. Brain structure images, after bias field correction and brain extraction (i.e., files of type *T1w_acpc_dc_restore_brain.nii.gz*) [63] were used for each subject. The strength of hand preference had been assessed with the Edinburgh Handedness Inventory [64], resulting in scores ranging from - 100 (strong left-hand preference) to 100 (strong right-hand preference). In addition, 30 HCP subjects had been scanned twice, so that test-retest analysis was possible in these 30 subjects (age ranges from 22-35 years at the scanning time; 20 females, 10 males).

#### BIL&GIN

BIL&GIN ([65]; *N* = 453; 232 females, age ranges 18-57 years at the scanning time). A high resolution T1-weighted MRI image was used for each individual, and brain images after bias field correction and brain extraction (implemented in *FreeSurfer* v5.3; surfer.nmr.mgh.harvard.edu) were used. Unlike the UK Biobank and HCP, which had natural population proportions of right-handers, BIL&GIN participants had been selected to be roughly balanced for handedness (248 right-handers and 205 left-handers), based on responses to the options: “*right-handed, left-handed, or forced left-handed*”. The strength of hand preference had again been assessed with the Edinburgh Handedness Inventory [64], resulting in scores ranging from −100 (strong left-hand preference) to 100 (strong right-hand preference).

### Global Asymmetry Measurement from T1-weighted Brain Images

A registration-based approach was used for global asymmetry measurement (Fig. 1), similar to that previously used in a study of vervet monkeys [12]. Specifically, for each individual participant, an affine transformation was applied to align the T1-weighted brain image (in native space) to the target template image (in the standard MNI space), and an affine transformation matrix was generated as an output. Image processing tools of *flirt* and *avscale* from FSL (version 5.0.10; fsl.fmrib.ox.ac.uk) were used for this analysis. The transformation matrix captures information about global shape differences between individual brain images and the target image, including scaling and skewing with respect to each axis. While the scaling factors are related to individual brain size, the skewing factors indicate the amount of global twisting to match the template. In order to measure left-right asymmetries, a left-right symmetrized template was used (i.e., ICBM 2009c Nonlinear Symmetric template). Here, we focused on skewing in the transverse (horizontal) and coronal (vertical) planes, to measure the two global asymmetry components, i.e. with respect to the frontal-occipital and dorsal-ventral axis, respectively (Fig. 1). A positive horizontal skew is closely akin to typical torque, i.e. the protrusions of the right frontal and left occipital regions, which has been reported as the average asymmetry pattern in the human brain (see Introduction). In contrast, a negative horizontal skew indicates a reversal of the typical pattern, with the left frontal and right occipital regions protruding. Similarly, a positive vertical skew indicates an overall twisting downward of the left hemisphere and upward of the right hemisphere, while a negative vertical skew indicates the opposite pattern.

**Fig. 1.**
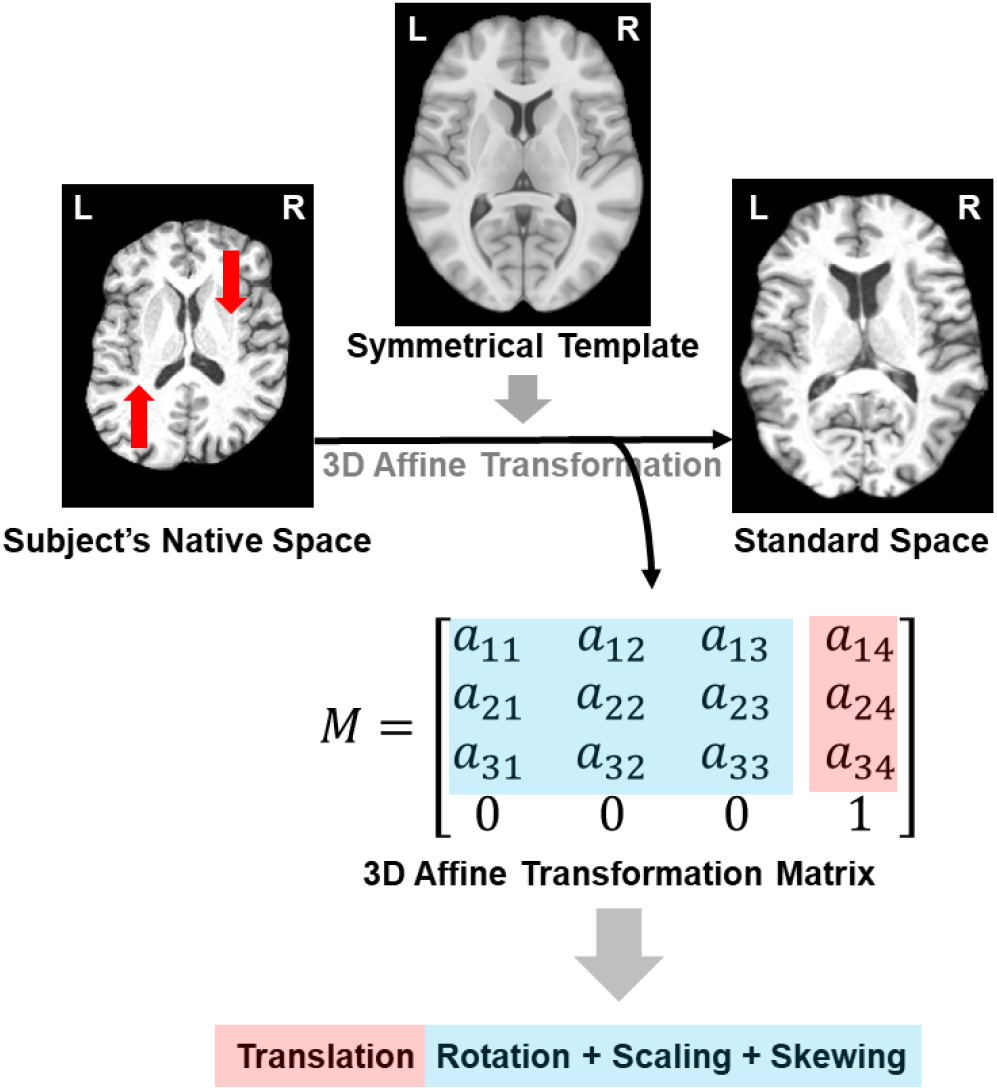
A registration-based approach for estimating global asymmetry skewing. The transformation matrix from the registration procedure captures information about the position alignment (i.e., translation and rotation) as well as scaling, and the amount of skewing during registration. Red arrows indicate the direction in which the native space image is shifted during image registration. Transverse sections are shown, which illustrate the horizontal skew process.

### Test-Retest Reliability of Global Asymmetry Measures

The HCP-Retest data (30 subjects with two scans per subject) allowed us to quantify test-retest reliability of the two global asymmetry metrics. Intraclass correlation coefficients (ICC) were calculated using IBM SPSS 20 (Model: Two-Way Mixed; Type: Consistency), where the ICC is conceptualized as the ratio of between-subjects variance to total variance. The ICC is a value between 0 and 1, where 1 indicates perfect reliability (i.e., the within-subject variance is 0).

### Association of Global Brain Asymmetry with Handedness

Association analyses of the horizontal and vertical skew measures were performed for each dataset separately, as per the availability of specific handedness measures and co-variables. In the UK Biobank, the asymmetry differences between handedness groups (−1=left, 0=both, 1=right, treated as a continuous variable) were assessed with linear regression models adjusted for sex, age, nonlinear age (z-transformed square of the age, zage^2^), the first ten principal components (PCs) which capture genome-wide population structure in the genotype data (as made available by the UK Biobank (PC1-10)) [62], and several technical variables related to imaging [61]: imaging assessment center (binary), scanner position parameters (continuous X/Y/Z), and signal/contrast-to-noise ratio in T1 (continuous). To explore which handedness group differences mainly contributed to the results, we repeated the analyses with each pair of handedness groups (i.e., right-handers vs. left-handers; right-handers vs. both-handers; and both-handers vs. left-handers). To exclude possible outliers in asymmetry measures, we excluded subjects above/below 4 standard deviations from the mean, separately for the horizontal and vertical skew measures. In addition, analyses were repeated when additionally adjusting for brain volume (i.e., grey+white volume) and the brain-size related scaling factors indicated in Figure 1. Python’s *pandas* (pandas.pydata.org) and *statsmodels* (www.statsmodels.org) packages were used for these analyses.

In the HCP dataset, asymmetry differences related to the strength of hand preference (ranging from −100 to +100, see above) were examined with linear regression models, adjusting for sex, age, nonlinear age (zage^2^), ‘*Acquisition*’ (a variable to control for possible scanner status differences across the study period of several years), and ‘Race’ (a variable given this label in the HCP data, to capture ethnicity). Analyses were repeated when additionally adjusting for brain volume (‘*FS_IntraCranial_Vol*’, intracranial volume estimated with FreeSurfer,) and the brain-size related scaling factors indicated in Figure 1. The HCP dataset included twins and siblings, and we therefore used Permutation Analysis of Linear Models (PALM, version alpha111) [66] from FSL (version 5.0.10), which has a specialized function for accounting for possible non-independence caused by family structure. For this we used 10,000 permutations and calculated 2-tailed *p* values.

In the BIL&GIN dataset, asymmetry differences related to the strength of hand preference (again ranging from −100 to +100, see above) were examined with linear regression models, adjusting for sex, age, and nonlinear age (zage^2^). Analyses were repeated when additionally adjusting for brain volume (again intracranial volume estimated with FreeSurfer) and the brain-size related scaling factors indicated in Figure 1. Python’s *pandas* and *statsmodels* packages were used. Although left-handedness was deliberately over-represented in the BIL&GIN dataset (to achieve balance for handedness), we did not attempt to correct for sampling bias, because the findings (see Results) were in line with the other two datasets, in which handedness was not over-represented. The relatively small sample size of the BIL&GIN dataset meant that repeat dropping of 9/10 of left handers to match their population prevalence would make the statistical power too low, and would also result in unequal group sizes (which can create its own statistical issues).

### Phenome-wide Associations of Global Brain Asymmetry

The UK Biobank dataset includes many variables, including early life factors, psychosocial factors, derived imaging traits, and variables related to cognitive functions and health. We ran Phenome-wide Association Scan (pheWAS) analysis for each global asymmetry component, to screen for other associated variables besides handedness and genetic data. We used the package PHEnome Scan Analysis Tool (PHESANT) [88], which enables comprehensive phenome scans to be performed across all data fields in the UK Biobank. PHESANT uses a rule-based method to automatically determine how to test each variable. The decision rules start by assigning each variable as one of four types: continuous, ordered categorical, unordered categorical, or binary. A description of PHESANT’s automated rule-based method is given in detail elsewhere [88]. PHESANT then estimates the bivariate association of an independent variable of interest (in our case either horizontal or vertical brain asymmetry) with each dependent variable in the dataset. Dependent variables with continuous, binary, ordered categorical, and unordered categorical data types, are tested using linear, logistic, ordered logistic, and multinominal logistic regression, respectively. Prior to testing, an inverse normal rank transform is applied to variables of the continuous data type. All analyses were adjusted for covariates as in the handedness association analyses (see above).

We corrected for multiple testing using Bonferroni correction, with a significance threshold determined by dividing 0.05 by the number of tests performed, separately for the horizontal and vertical asymmetry measures as independent variables. We also looked up the results for some variables of particular interest in relation to global brain asymmetry (see Results), which did not necessarily survive multiple testing correction over all phenotypes tested, in which case we report the nominal P values. This enables comparison of the identified associations with results for other, possibly related variables.

Finally, as language is a prominently lateralized and human-specific function [89, 90], we ran association analyses between the two global asymmetry components and four behavioral performance measures related to language, which were available in the HCP dataset. The tasks included were the Penn Word Memory Test, Language Task for fMRI, and the NIH Toolbox Oral Reading Recognition Test and Picture Vocabulary Test [91]. PALM was again used for accounting for family structure in the statistical analyses (see above). Multiple testing was corrected using the Bonferroni method (corrected *p* < 0.05).

### Heritability and Polygenicity Estimation

In the UK Biobank, 550,192 autosomal, directly genotyped SNPs with minor allele frequencies (MAF) > 0.01, genotyping rate >0.95 and Hardy-Weinberg equilibrium (HWE) p>1×10^−6^ were used to build a genetic relationship matrix (GRM) using GCTA (version 1.26.0) [67]. We excluded samples with a genotyping rate of <98% and a kinship coefficient higher than 0.025 based on this GRM, resulting in a sample size of 30,682. Genome-based restricted maximum likelihood (GREML) analyses using GCTA were performed to estimate the SNP-heritabilities for the horizontal and vertical skew measures, after residualizing for the covariate effects of sex, age, zage^2^, the first ten principal components capturing genome-wide genetic structure [62], genotyping array, and several technical variables related to imaging as mentioned above. SNP-based heritability is a measure ranging from 0 to 1 which indicates the extent to which variation in a trait is influenced by the combined effects of variations at SNPs distributed over the genome [68]. Bivariate analyses [69] were also run in GCTA, to investigate the SNP-based genetic correlations between the two global asymmetry measures, and also with the x, y and z scaling factors that indicate brain size (see above). Genetic correlation analysis measures the extent to which variability in a pair of traits is influenced by the same genetic variations over the genome.

In addition, we also estimated SNP-based heritability of the two global asymmetry measures in the UK Biobank data using GWAS summary statistics for each measure (see below), using LD-score regression as implemented in the LDSC package (v1.0.1) (https://github.com/bulik/ldsc) [70]. This approach was also used to measure genetic correlation between brain skews and left-handedness, using GWAS summary statistics for left-handedness reported by de Kovel et al [53]. Precomputed LD scores from the 1000 Genomes European data (i.e., *eur_w_ld_chr.tar.bz2*) were used, and there was no constraint on the intercept of regression. We also applied causal mixture models to estimate polygenicity (estimated number of causal variants) and discoverability (proportion of phenotypic variance explained on average by a causal variant, σ_β_^2^) [71], using the MiXeR package (v1.2) (https://github.com/precimed/mixer). This analysis was based on the GWAS summary statistics for the brain skew measures as generated in the present study.

Heritability could also be estimated in the HCP dataset, as it included monozygotic and dizygotic twin pairs, as well as unrelated individuals (see above). We estimated the heritability of each global asymmetry component using variance-component analysis implemented in SOLAR [72]. Briefly, each asymmetry component was entered as a dependent variable into separate linear mixed-effects models, which included fixed effects of sex, age, and nonlinear age (zage^2^), and a random effect of genetic similarity, whose covariance structure was determined by the pedigrees. Genetic similarity for MZ pairs is coded as 1 as they share 100% of their DNA sequence, whereas DZ twins and siblings are coded as 0.5 as they share on average 50%, while unrelated individuals are coded as zero. Maximum likelihood-based bivariate variance decomposition analysis was also applied, again using SOLAR, to estimate the genetic correlation of the two global asymmetry measures.

### Genome-wide Association Scans

Imputed SNP genotype data (bgen files; imputed data v3 – release March 2018) were extracted for the samples with global brain asymmetry measures (*N* =33,996), and SNP-level statistics were then computed within this set using QCtools (v.2.0.1). We excluded SNPs with a minor allele frequency below 1%, Hardy-Weinberg p-value below 1×10^−7^ or imputation quality INFO scores below 0.7 (the latter as provided by the UK Biobank with the imputed data [62]), which resulted in 9,904,141 SNPs genome-wide. GWAS was performed with BGENIE (v.1.2) [73] for each of the residualized global asymmetry measures separately (after accounting for the same covariate effects as for SNP heritability analysis, above), using imputed genotype dosages and an additive model. We applied the commonly-used genome-wide significance threshold *p* value of 5e-08 to assign significance in the context of genome-wide multiple testing, which accounts for the number of SNPs tested in a modern GWAS study, and the correlation structure between SNPs in European ancestry populations [74, 75].

### Gene-based and Gene-set Analyses

We derived gene-level association statistics based on the GWAS summary statistics using MAGMA (v1.08) [76] implemented in FUMA (v1.3.6) [77]. In brief, the gene-wise test summarizes the degree of association between a phenotype and SNPs within a given gene [76]. The gene window was set to 50 kb upstream and downstream to include nearby cis regulatory regions. European samples from the 1000 Genomes phase 3 were used as a reference panel to account for linkage disequilibrium (LD) between SNPs. A significance threshold of *p* < 2.481e-06 (i.e., 0.05/20151) was applied to correct for multiple testing across all protein coding genes (Ensembl version v92; n = 20,151), and a further Bonferroni correction was also considered for having studied two skew measures.

The gene-level association statistics were then used to perform gene-set enrichment analysis, again using MAGMA, for gene ontology (GO) terms for biological processes, cellular components, and molecular functions [78] (minimum set size of 10 genes, maximum size 1000 genes, total *n* = 6576 GO sets meeting these criteria) obtained from the latest MsigDB (v6.2) database (http://software.broadinstitute.org/gsea/msigdb), and Bonferroni correction was performed. This approach tests whether the genes in a given set show, on average, more evidence for association with the trait in question than the rest of the genes in the genome for which scores could be calculated, while accounting for non-independence of SNPs due to LD.

### Genetic Correlation with Psychiatric Disorders

We calculated genetic correlations of the horizontal and vertical skew measures with three psychiatric disorders that have been prominently reported to associate with altered brain asymmetry: ASD [79], ADHD [20], and SCZ [49, 80]. This analysis was based on GWAS summary statistics for the brain skew measures as generated in this study, together with publicly available GWAS summary statistics for ASD (18,381 cases and 27,969 controls) [81], SCZ (36,989 cases and 113,075 controls)[82], and ADHD (20,183 cases and 35,191 controls)[83]. We used the publicly available summary statistics of the largest-to-date GWAS for each of these disorders. The LDSC package (https://github.com/bulik/ldsc) [70] was used for calculating genetic correlations, and Bonferroni correction was performed for three disorders and two skews.

## Results

### Global Brain Asymmetry in the UK Biobank, HCP, and BIL&GIN datasets

We extracted two global asymmetry components for each individual. A positive score for horizontal skew indicates a global pattern in which the right hemisphere is shifted anteriorly relative to the left, while a negative score indicates that the left hemisphere is shifted anteriorly relative to the right (see Fig. 2 and Supp. Fig. 1 for examples). Note that the skew is calculated as a global feature, not a feature of any particular slice (Supp. Fig. 1). Similarly, a positive score for vertical skew indicates a global shift downwards of the left hemisphere relative to the right, while a negative vertical skew indicates the opposite pattern (Fig. 2, Supp. Fig. 2). Both asymmetry components showed almost perfect reliability (horizontal skew: *ICC* = 0.989; vertical skew: *ICC* = 0.977) as indicated in test-retest analysis of data from 30 HCP subjects who underwent two scans each (Fig. 2).

**Fig. 2.**
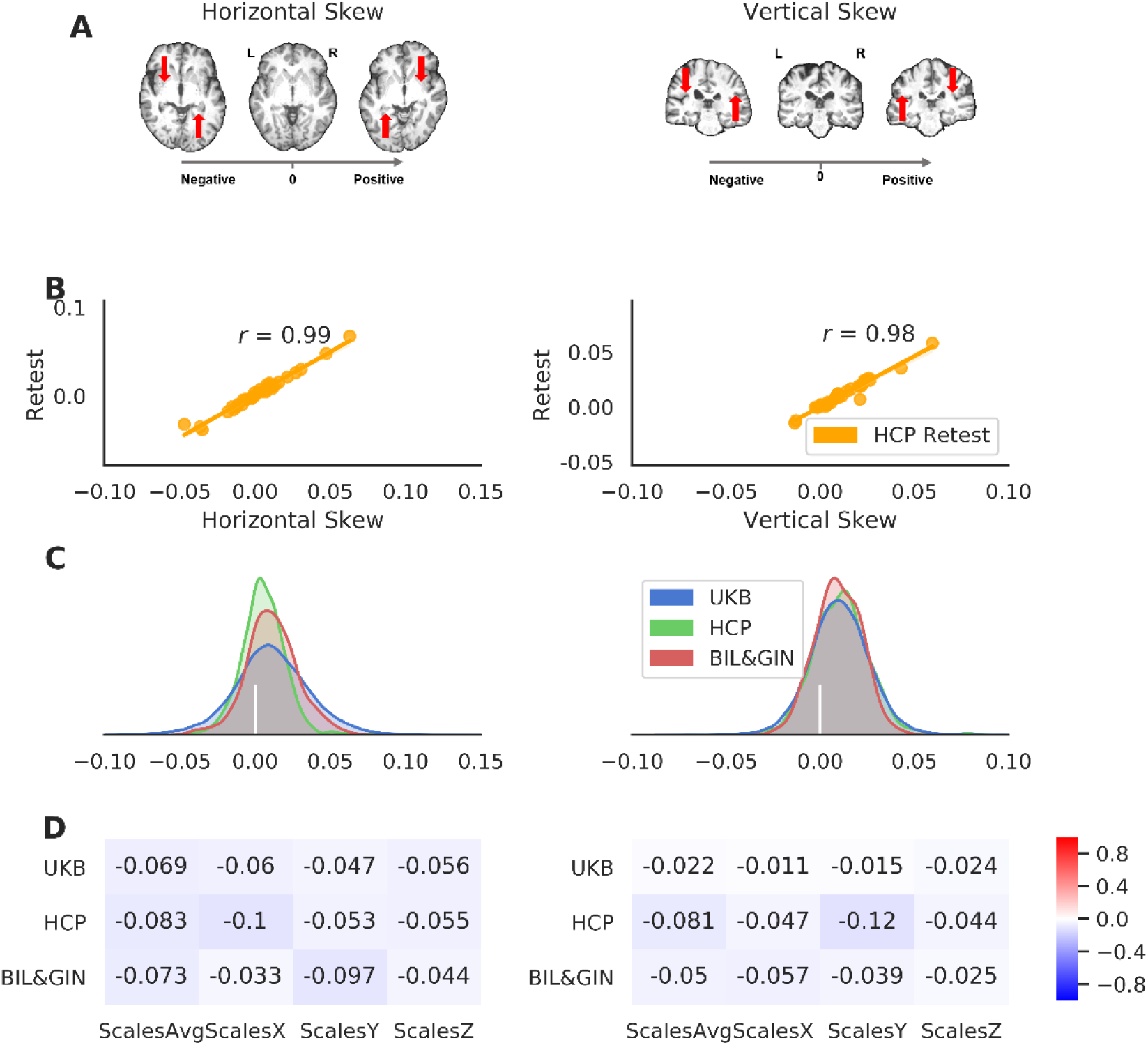
Global brain asymmetry: the horizontal and vertical skews. (A) Examples of the human brain with different asymmetry skew scores. The right-pointing grey arrow indicates the axis of the skew scores from negative to positive. The red arrows indicate the skewing needed for each brain during image registration (i.e., registration of the native space brain to a symmetrical template). (B) Scatter plots of the asymmetry skews in the HCP-Retest dataset, with the Pearson correlation coefficients. (C) Distributions of the asymmetry skew scores in the three datasets: UK Biobank in blue, HCP in green, and BIL&GIN in red. The vertical bar in white indicates the position of zero skewing. (D) The Pearson correlation coefficients of the asymmetry skew scores with brain size-related scaling factors in the three-dimensions (ScalesX/ScalesY/ScalesZ) and their average (ScalesAvg).

The average values of both asymmetry components were positive in the three independent datasets (Fig. 2), confirming a population-level pattern of global asymmetry. For horizontal skew, the average pattern involved protrusions of the right frontal and left occipital regions (UK Biobank: *N* = 39,678, *Mean* =0.0110, *Std* = 0.025; HCP: *N* = 1113, *Mean* = 0.0054, *Std* = 0.014; BIL&GIN: *N* = 453, *Mean* = 0.011, *Std* = 0.017). One sample t-testing indicated a significant difference of each dataset mean from zero (UK Biobank: *t*(39,677) = 86.78, *p* < 5.00e-100; HCP: *t*(1112) = 12.56, *p* = 5.96e-34; BIL&GIN: *t*(452) = 14.14, *p* = 7.68e-38). The population average horizontal skew matches the widely-observed features of brain torque (e.g., frontal/occipital petalia) in the human brain (See Introduction and e.g. Thompson et al., 2003).

Regarding the vertical asymmetry skew, the sample means were again all positive (UK Biobank: *Mean* = 0.0103, *Std* = 0.0158; HCP: *Mean* = 0.011, *Std* = 0.015; BIL&GIN: *Mean* = 0.0097, *Std* = 0.012), indicating an average pattern involving downward skewing of the left hemisphere relative to the right. Again the means were significantly different from zero in each dataset (UK Biobank: *t*(39,677) = 131.06, *p* < 5.00e-100; HCP: *t*(1112) = 24.48, *, p < 5.00e-100*; BIL&GIN: *t*(452) = 16.96, *p* = 2.90e-50). Notwithstanding the average asymmetry patterns, the distributions of the vertical and horizontal skews showed considerable individual differences, with e.g. 31.8% of participants showing a reversal compared to the average horizontal pattern, and 24.6% showing a reversal compared to the average vertical pattern, in the UK Biobank dataset (Fig. 2). The horizontal and vertical skews showed low correlations with brain-size-related scaling factors in the three datasets (Fig. 2): UK Biobank (|*r*|s < 0.07), HCP (|*r*|s < 0.12), and BIL&GIN (|*r*|s < 0.097) (Table S1), such that the skews appear to be largely independent of brain size. Also, the two skews showed low correlations with each other, and these were inconsistent in strength and direction among the three datasets: UK Biobank (*r* = 0.074), HCP (*r* = −0.170), BIL&GIN (*r* = −0.030).

### Global Brain Asymmetry and Handedness

We found significant associations between handedness (left coded as −1, *N* = 3501; ambidextrous coded as 0, *N* = 572; right coded as 1, *N* = 35338) and both asymmetry components in the UK Biobank (Fig. 3; horizontal skew: t = 5.15, p = 2.56e-07; vertical skew: t = −9.18, p = 4.36e-20). These associations were mainly contributed by group differences between left-handers and right-handers (horizontal skew: *t* = 5.08, *p* = 3.77e-07, *d* = 0.090; vertical skew: *t* = −9.31, *p* = 1.34e-20; *d* = 0.165). These results show that right-handers, compared with left-handers, are more likely to show skew along the anterior-posterior axis in the same direction as the population average pattern (i.e., more positive horizontal skew scores). However, on the dorsal-ventral axis, left-handers are more likely to show skew in the same direction as the population average pattern (i.e., more positive vertical skew scores). In addition, a difference was found when comparing vertical skew between left-handers and ambidextrous participants (t = −3.10, p = 0.00195), while no other comparisons between handedness groups showed significant effects (*p*s > 0.15).

**Fig. 3.**
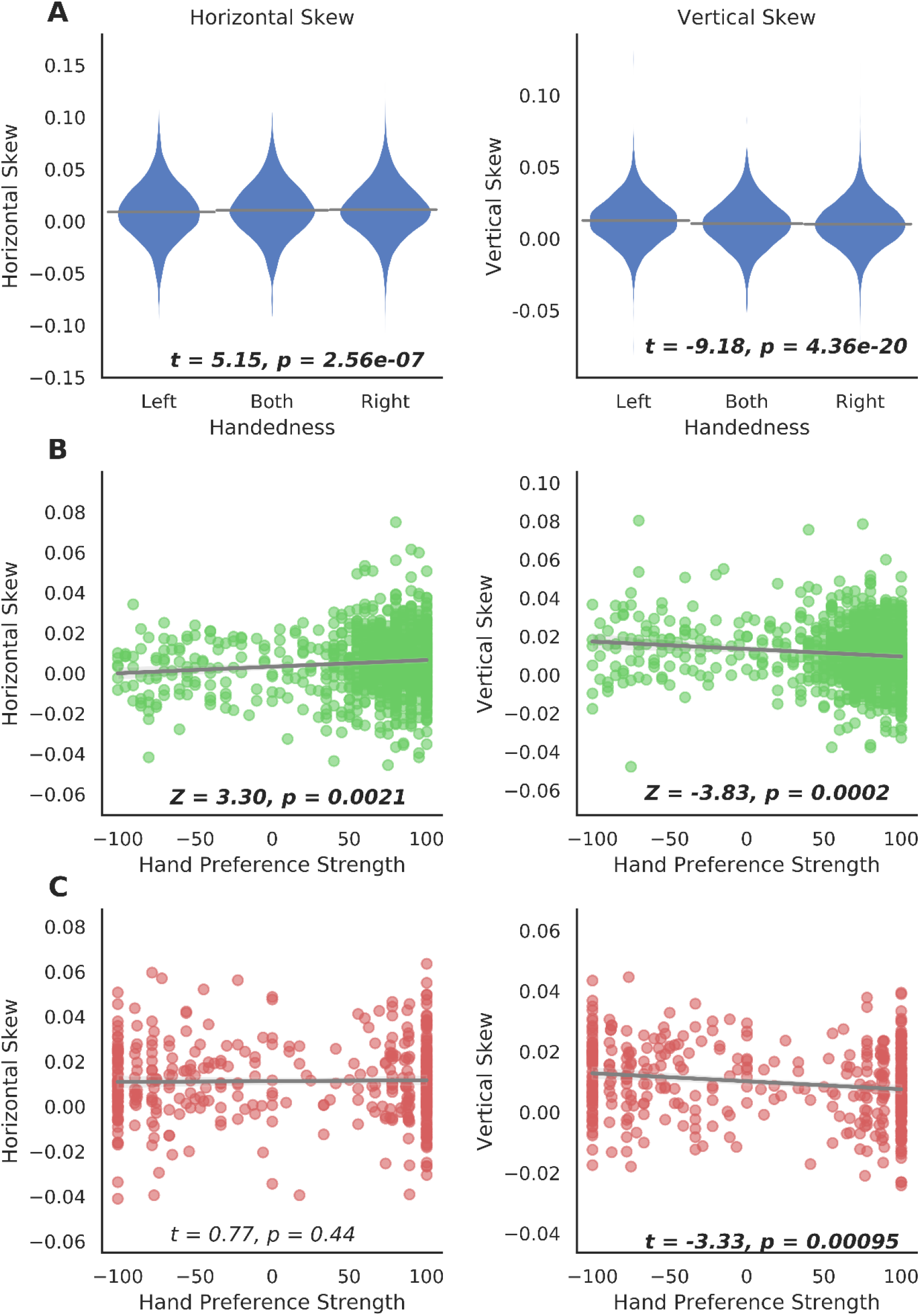
Global asymmetry skews and hand preference. (A) Differences in global asymmetry measures between handedness groups in the UK Biobank. (B) Scatter plots of global asymmetry measures and hand preference strength in the HCP. (C) Scatter plots of global asymmetry measures and hand preference strength in BIL&GIN. Note that the statistical tests of association were based on analyses with covariate effects being controlled for, whereas data are plotted here without adjusting for covariates, for display purposes.

The HCP dataset confirmed that the two skews were related to hand preference. Here, a continuous index of the strength of hand preference was available, from −100 (left) to 100 (right), and the analysis accounted for family structure (see Methods). Again, horizontal skew was positively associated with right-hand preference (*Z* = 3.30, *p* = 0.0021), and vertical skew was positively associated with left-hand preference (*Z* = −3.83, *p* = 0.0002) (Fig. 3).

The BIL&GIN dataset further confirmed the finding with respect to vertical skew, i.e. this was again positively correlated with increased left-hand preference, *t* = −3.33, *p* = 0.00095 when using the hand preference scale from −100 to 100, and *t* = −3.61, *p* = 0.00035 when using a binary handedness assessment, see Methods (Left: *N* = 205; Right: *N* = 248). However, in this dataset, the association of hand preference with horizontal skew was not significant, *t* = 0.77, *p* = 0.44 for the continuous hand preference scale, and *t* = 0.81, *p* = 0.42 for binary handedness. Nonetheless, the direction of this non-significant association in BIL&GIN was consistent with the UK Biobank and HCP datasets. The BIL&GIN dataset provided only 24.4% power to detect the association of horizontal skew with handedness at alpha 0.05, according to the effect size of this association in the UK Biobank (Cohen’s *d* = 0.090). The non-significant association between horizontal skew and handedness in BIL&GIN is therefore likely to be a power issue, due to a relatively limited sample size (whereas for the association with vertical skew, BIL&GIN provided 54% power at alpha 0.05, in relation the UK Biobank effect size *d* = 0.165).

When adjusting vertical skew for horizontal skew, and vice versa, the associations of both asymmetry variables with hand preference remained very similar (UK Biobank, three handedness groups, horizontal skew, *t* = 5.70, *p* = 1.23e-08; vertical skew, *t* = −9.50, *p* = 2.23e-21: HCP, horizontal skew, *Z* = 2.69, *p* = 0.011; vertical skew, *Z* = −3.32, *p* = 0.0006: BIL&GIN, hand preference strength, horizontal skew, *t* = 0.67, *p* = 0.50; vertical skew, *t* = −3.57, *p* = 0.00039). This indicates that handedness is primarily associated independently with the two asymmetry variables, rather than with any shared variance between them.

In the UK Biobank, there was a significant association between handedness (including left, ambidextrous and right groups) and one of the brain-size-related scaling factors (i.e., ScalesY: scaling in the anterior-posterior direction, *t* = −3.12, *p* = 0.00174). However, this association did not replicate in the HCP (*t*s<1) or BIL&GIN datasets (*t*s<1 except for ScalesZ: scaling in the superior-inferior direction, *t* = 1.75, *p* = 0.08). Moreover, when controlling for these scaling factors related to brain size, associations between hand preference and the global asymmetries remained unchanged (UK Biobank, three handedness groups, horizontal skew, *t* = 5.17, *p* = 2.32e-07; vertical skew, *t* = −9.21, *p* = 3.57e-20; HCP, horizontal skew, *Z* = 3.38, *p* = 0.0014; vertical skew, *Z* = −3.60, *p* = 0.0004; BIL&GIN, hand preference strength, horizontal skew, *t* = 0.89, *p* = 0.38; vertical skew, *t* = −3.38, *p* = 0.00079).

#### Phenome-wide associations of brain skew measures

In the UK Biobank dataset, we ran phenome-wide association screening (pheWAS) to search for other variables associated with the horizontal or vertical components of global brain asymmetry. The variables were those included in our approved UK Biobank project 16066 ‘Genetics of brain asymmetry and language-related disorders’, and consisted mainly of measures related to cognitive functions, sociodemographics, mental health, physical measures, medical information, and brain imaging measures. In total, the pheWAS analysis included 3562 tests for each of the two skew measures. Note that some variables might only be considered as ‘phenotypes’ in a very broad sense, such as country of birth, and home area population density.

For each of the two skew measures, the pheWAS QQ plot is shown in Fig. S3. There were 464 associations below the Bonferroni corrected threshold of 1.40e-05 (0.05/3562) for horizontal skew, and 293 associations for vertical skew. Horizontal skew showed significant associations with variables of various categories, including cognitive functions (e.g., “*Fluid intelligence score*”: *p* = 2.74e-06), sociodemographics (e.g., “*Age completed full time education*”: *p* = 6.19e-08), physical measures (e.g., “*Body mass index*”: *p* = 6.05e-07), and mental health (“*Recent changes in speed/amount of moving or speaking*” from the *Depression* test: *p* < 1.00e-155) (Fig. 4A and Dataset S1). The overwhelming majority (440 out of 464) were associations with brain imaging variables (Fig. 4). Interestingly, these latter associations showed a global anterior-posterior “torque” pattern for both grey matter volume (Fig. 4B) and white matter metrics (Fig. 4C). In terms of grey matter, the volumes of right frontal and left occipital regions positively correlated with horizontal skew, while left frontal and right occipital regional volumes negatively correlated with horizontal skew (Fig. 4B). Similarly, microstructural measures of homologous white matter tracts showed significant associations with skew measures in opposite directions in the two hemispheres (Fig. 4C). This confirms that torque manifests not only as a relative shifting of the hemispheres overall, but also as interhemispheric regional asymmetries affecting both grey and white matter [6]. More details can be seen in Fig. S5A, Fig. S6, and Dataset S1.

**Fig. 4.**
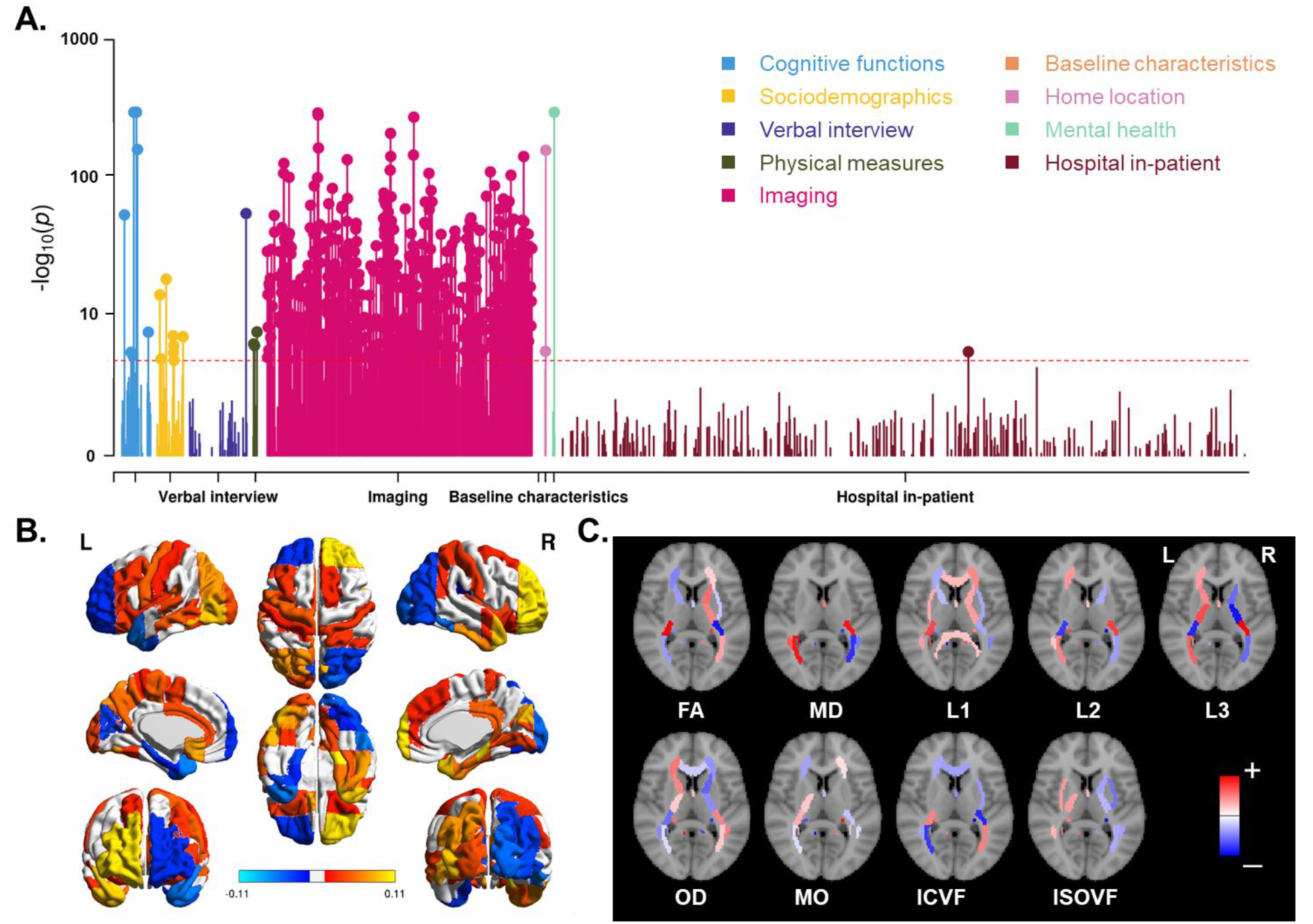
Phenome-wide association analysis for horizontal brain skew. (A) Manhattan Plots for the associations. Red lines indicate the Bonferroni corrected threshold (p < 1.40e-05). (B) Significant associations of skew measures with regional grey matter volumes. Red-yellow indicates a positive association; blue indicates a negative association. (C) Significant associations of skews with various white matter metrics. Red indicates a positive association; blue indicates a negative association. The per-region names and statistics for parts B and C can be found in Dataset S1. FA: fractional anisotropy, MD: mean diffusivity, L1/ L2/L3: the three eigenvalues of diffusion, MO: mode of anisotropy, OD: orientation dispersion, ICVF: intra-axonal volume fraction, ISOVF: isotropic volume fraction.

Regarding the vertical skew, we found significant associations with variables of various categories, including cognitive functions (e.g., “*Time to answer*”/Prospective memory: *p* = 1.91e-08), sociodemographics (e.g., “*Transport type for commuting to job workplace: Car/motor vehicle*”: *p* = 6.19e-08), physical measures (e.g., “*Body mass index*”: *p* = 1.20e-05), and mental health (“*Recent changes in speed/amount of moving or speaking*” from the *Depression* test: *p* < 4.66e-77) (Fig. 5A and Dataset S2). Again these significant associations were mostly (274 out of 293) with brain imaging variables. As is shown in Fig. 5B, grey matter volumes of the left inferior, medial temporal and occipital regions correlated positively with vertical skew, while the homologous regions in the right hemisphere showed negative correlations. The microstructural metrics of white matter tracts also showed a similar complementary interhemispheric pattern (Fig. 5C). More details can be seen in Fig. S5B, Fig. S7, and Dataset S2.

**Fig. 5.**
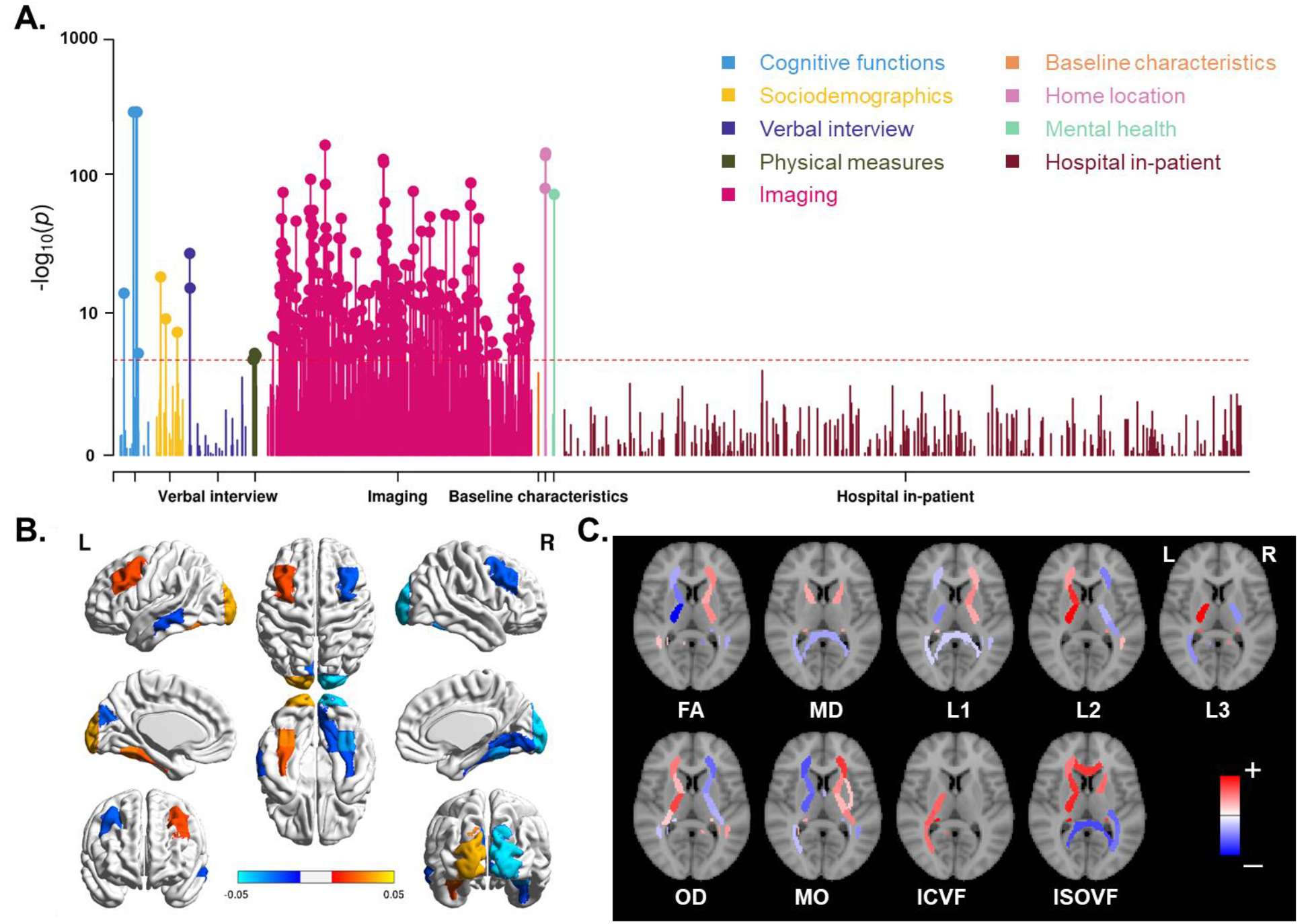
Phenome-wide association analysis for vertical brain skew. (A) Manhattan Plots for the associations. Red lines indicate the Bonferroni corrected threshold (p < 1.40e-05). (B) Significant associations of skew measures with regional grey matter volumes. Red-yellow indicates a positive association; blue indicates a negative association. (C) Significant associations of skews with various white matter metrics. Red indicates a positive association; blue indicates a negative association. The per-region names and statistics for parts B and C can be found in Dataset S2. FA: fractional anisotropy, MD: mean diffusivity, L1/ L2/L3: the three eigenvalues of diffusion, MO: mode of anisotropy, OD: orientation dispersion, ICVF: intra-axonal volume fraction, ISOVF: isotropic volume fraction.

Certain early life factors have been shown to influence handedness in the UK Biobank, including birthweight, multiple birth, breastfeeding, and country of birth within the UK [29]. Maternal smoking was not found to influence handedness in the UK Biobank [29], although has been implicated in handedness by other studies (e.g. [84]). We looked specifically into the pheWAS results for these early life factors, for the two brain skew measures. Both skew measures showed significant associations with variables related to place of birth (“*Country of birth*”, *p* = 3.53e-15 for horizontal skew, Dataset S1; “*Place of birth in UK - north co-ordinate*”, *p* = 2.56e-29 and “*Place of birth in UK - east co-ordinate*”, *p* = 8.32e-17 for vertical skew, Dataset S2). In addition, breastfeeding (*p* = 0.0030) and birth weight (*p* = 0.025) showed nominally significant associations with vertical skew, while other associations were not significant (*p*s > 0.10).

In the HCP dataset, analysis of four behavioral performance measures related to language (see Methods) showed a correlation between horizontal skew and oral reading recognition performance (*Z* = 3.37, permuted *p* = 0.0023, which survived Bonferroni correction for eight tests, i.e. two skew measures times four performance measures), such that individuals with more positive horizontal skew showed better oral reading recognition ability. We found no associations with other behavioral measures in the HCP (unadjusted *p*s > 0.10).

### Heritability and gene mapping for brain skews

In the UK Biobank, significant but low SNP-based heritabilities were found for horizontal skew (*h*^*2*^ = 3.85%, *95% confidence interval (CI)* = [0.70%, 7.00%], *p* = 0.024 using GCTA [67]; *h*^*2*^ = 4.67%, *95% CI* = [0.63%, 8.89%], *p* = 0.029 using LDSC [70]) and vertical skew (*h*^*2*^ = 7.64%, *95% CI* = [4.42%, 10.86%], *p* = 3.01e-05 using GCTA [67], *h*^*2*^ = 12.95%, *95% CI* = [8.94%, 16.96%], *p* = 5.56e-08 using LDSC [70]). In the HCP dataset, which included twins, both skew measures also showed evidence for low heritability: horizontal skew *h*^*2*^ = 9.1%, *p* = 0.041; vertical skew *h*^*2*^ = 10.1%, *p* = 0.030. Vertical skew was slightly more heritable than horizontal skew in both datasets. These findings suggest that genetic variability influences global brain asymmetry, but that most of the variance is not due to genetic variation.

In the UK Biobank data, we also applied causal mixture models [71] to estimate the polygenicity (estimated number of causal variants) and discoverability (proportion of phenotypic variance explained on average by each causal variant, σ_β_^2^) for each skew measure. The results indicated that vertical skew has a higher polygenicity (5377 causal variants, AIC = 1.80, BIC = 12.51) than horizontal skew (1111 causal variants, AIC = 1.92, BIC = 12.64), at a similar level of discoverability (σ_β_^2^ = 1.47e-05 vs. σ_β_^2^ = 2.94e-05).

Genome-wide association analyses showed no significant loci for either horizontal or vertical skew with the standard genome-wide significance threshold of 5e-08 [74, 75] (Fig. S4), consistent with low heritability and substantial polygenicity. Five SNPs with suggestive association p values less than 5e-07 are listed in Table S2 (one SNP for horizontal skew and four for vertical skew), together with their association statistics and nearest genes. Similarly, in gene-based association analysis [76], no significant genes were found for either horizontal or vertical skew at the genome-wide, gene-based significance threshold *p* < 2.48e-06 (i.e., 0.05/20151 protein coding genes). The most significant genes were *PLB1* for horizontal skew (chr.2, *Z* = 3.93, *p* = 4.20e-5) and *PLEC* (chr.8, *Z* = 4.34, *p* = 7.12e-6), *GRINA* (chr.8, *Z* = 4.24, *p* = 1.11e-5) and *PARP10* (chr.8, *Z* = 4.08, *p* = 2.26e-5) for vertical skew.

Gene-set analysis of the GWAS results for vertical skew, using MAGMA [76], showed a significant enrichment of association within the Gene Ontology term “*BP:go_neuron_projection_guidance*” (*beta* = 0.21, *p* = 7.59e-6) after correction for multiple testing (*p* < 7.60e-06, i.e., 0.05/6576 gene sets), which would not be significant with further correction for two skews tested. There were no significant sets identified for horizontal skew. Top gene sets with nominal P values < 0.001 are listed in Table S3.

### Genetic correlations of brain skews with other traits

Handedness has shown a low but significant heritability in the UK Biobank in two previous studies based on more than 330,000 individuals: *h*^2^ =1.8%, 95% CI = [1.79%, 1.81%] [53]; *h*^2^ = 1.2%, *95% CI* = [1.197%, 1.202%] [29, 52] (note that the sample sizes for those studies were much larger than the present study, because the present study is limited to participants who also have brain MRI data available). However, using LD score regression with the GWAS summary statistics for handedness (*N* > 330,000) from de Kovel et al. [53] and the summary statistics from our GWAS for brain skew measures, we found no significant genetic correlations of hand preference with the skews (*p*s > 0.10). GCTA-based genetic correlation analysis within those individuals having both handedness and brain imaging data (*N* = 32,774) showed similar results (*p*s > 0.10). This was also the case using twin-based co-heritability analysis in the HCP dataset, i.e. no significant genetic correlations of handedness and brain skews (*p*s > 0.10).

Brain size-related scaling factors showed high heritabilities in both the UK Biobank (ScalesAvg: *h*^2^ = 72.11%; ScalesX: *h*^2^ = 56.35%; ScalesY: *h*^2^ = 46.28%; ScalesZ: *h*^2^ = 60.36%) and the HCP (ScalesAvg: *h*^2^ = 92.1%; ScalesX: *h*^2^ = 85.5%; ScalesY: *h*^2^ = 85.0%; ScalesZ: *h*^2^ = 86.6%), which was expected because of the high heritability of brain size (UK Biobank: *h*^2^ = 72.3%; HCP: *h*^2^ = 87.8%). There were no significant genetic correlations between these scaling factors and brain skew measures (*p*s > 0.10) (again using GCTA in the UK Biobank and twin-based co-heritability analysis in the HCP), except for a consistent negative genetic correlation between horizontal skew and ScalesX (scaling in the left-right axis) (UK Biobank: *r*_*g*_ = −0.212, *p* = 0.0343; HCP: *r*_*g*_ = −0.291, *p* = 0.0341, *p* values not adjusted for multiple testing). Thus some of the same genetic variability that contributes to a wider brain may also contribute to a more positive horizontal skew (e.g., anterior shift of the right hemisphere).

In addition, within the UK Biobank data, we ran genetic correlation analyses for brain skews in relation to traits that showed significant associations in the pheWAS analysis, using GCTA. Eight genetic correlations were nominally significant (*p* < 0.01), including grey matter volume in the left Heschl’s Gyrus (*Z* = 2.36, *p* = 0.0093) with horizontal skew, mean MO (mode of anisotropy) in the left uncinate fasciculus (*Z* = −2.40, *p* = 0.0083) with horizontal skew, and place of birth in UK - east co-ordinate (*Z* = −2.42, *p* = 0.0078) with vertical skew, but none survived multiple testing correction (Table S4). While horizontal skew showed a significant correlation with oral reading recognition in the HCP data (see above), no significant genetic correlation was observed between them (*p* = 0.54), using SOLAR [72].

Using the most recent GWAS summary statistics for three psychiatric disorders that have been proposed to associate with altered brain asymmetry [49, 79], i.e., autism spectrum disorder (ASD) [81], attention deficit/hyperctivity disorder (ADHD) [83], and schizophrenia (SCZ) [82], we found that horizontal skew showed evidence for genetic correlation (LDSC package [70]) with ASD (*r*_rg_ = −0.40, *p* = 0.0075, significant at *p*<0.05 after Bonferroni correction for three disorders and two skews), while the genetic correlation of ASD with vertical skew was weaker (*r*_rg_ = −0.17, *p* = 0.069). Other genetic correlations were not significant (*p*s > 0.10).

## Discussion

We carried out the largest-ever analysis of global brain shape asymmetry, i.e., the horizontal and vertical asymmetry skews, in three independent datasets. The largest of the datasets comprised over 39,000 participants. At the population level, there was an anterior and dorsal skew of the right hemisphere, relative to the left. The population variances of these skews were largely independent, but both showed replicable associations with handedness, which establishes a link between lateralized structure and function of the human brain. The two skews also showed associations with multiple regional grey and white matter metrics, as well as various phenotypic variables including cognitive functions, sociodemographic physical, and mental health measures. The two skews showed SNP-based heritabilities of 4-13% depending on the method used to assess this, but no significant loci were found in GWASs, probably due to substantial polygenicity together with relatively low heritability. There was evidence for a low but significant genetic correlation between horizontal skew and ASD, which may be consistent with the subtle but widespread alterations of cortical regional asymmetries observed in ASD in a recent large-scale study [52].

### Global Brain Asymmetry

We measured the anterior-posterior and superior-inferior aspects of global left-right asymmetry in the human brain, using three population datasets and an automated, registration-based approach. The approach contrasts with older, manual methods for evaluating global asymmetry, or approaches using regionally restricted frontal and/or occipital hemispheric differences as proxies for overall torque (e.g., [2, 21, 22, 85–87]. Rather, the registration-based approach of the present study allowed an automatic and objective assessment of global asymmetry based on skewing transformation of the brain as a whole. This provides a truly global measure of left-right asymmetry in the human brain. Moreover, the skew metrics in the present study showed high test-retest reliability in twice-scanned individuals.

We found population-level asymmetrical skews on both the horizontal and vertical axes. The average horizontal skew pattern was consistent with previously-observed features of brain torque in the human brain, involving a more anteriorly protruding right frontal lobe and posteriorly protruding left occipital lobe (i.e., frontal/occipital petalia), and relative increases in the dimensions (e.g., volume and width) of the right frontal and left occipital poles (see [7] for a review). The horizontal skew may also relate to a population-level, frontal-occipital asymmetry gradient in regional cortical thickness, recently reported in a large-scale study [6]. Indeed, we also observed in the present study that individual differences in horizontal skew showed positive correlations with grey matter volumes of right frontal and left occipital regions, and negative correlations with left frontal and right occipital regions, again in line with previously described features of brain torque [7].

The mean population-level asymmetry pattern in the vertical plane, i.e. along the inferior-superior axis, involved an overall twisting of the left hemisphere downward, and right hemisphere upward. This aspect of global brain asymmetry has not been described consistently in the literature, but we found it to be replicable in the three independent cohorts of this study. Our findings are also consistent with another recent report that the left-occipital pole is shifted significantly downward relative to the right, on average [8]. We found that individual differences in vertical skew correlated positively with grey matter volumes of left inferior temporal regions and the occipital pole, while correlating negatively with the homologous regions of the right hemisphere.

As both vertical and horizontal skews showed associations with numerous, widely-distributed regional grey and white matter metrics, then we can conclude that global asymmetry is not simply a spatial displacement of the left and right hemispheres with respect to one another. Rather, global asymmetry relates to structural differences between the two hemispheres, affecting many structures from front to back, and top to bottom, and thus likely relates to functionally meaningful hemispheric differences. Given that global asymmetry measures correlated with both regional grey matter volumes and white matter properties, our analysis does not support a previous suggestion that brain torque is driven by white but not grey matter [88].

An aspect that was inconsistent across datasets in the present study was the relationship between individual differences in the horizontal and vertical components of global asymmetry. Specifically, a positive correlation between horizontal and vertical skew of 0.074 was found in the UK Biobank, but a negative correlation of similar magnitude was observed in the HCP, and no significant correlation was present in the BIL&GIN. Given that the UK Biobank was by far the largest of the three datasets, a low, positive correlation between the vertical and horizontal skews is likely to be the true population pattern. Regardless, it is clear that the two components of global brain asymmetry are largely independent in their variabilities. Therefore, consideration of these two distinct aspects of global brain asymmetry will be important in future studies of their functional significances and phenotypic associations. In general, our results are consistent with other literature which indicate that multiple different asymmetries of the brain can vary largely independently of each other [6, 89–91].

### Global Brain Asymmetry and Handedness

We found significant associations of both horizontal and vertical skews with handedness, or the strength of hand preference. On average, left-handers showed relatively lower horizontal asymmetry scores than right-handers, i.e. reduced asymmetry along the anterior-posterior axis, and higher vertical asymmetry scores, i.e., increased asymmetry along the dorsal-ventral axis. The effect sizes were small, e.g. in the UK Biobank: Cohen’s *d* = 0.09 for the handedness association with horizontal skew, and *d* = 0.17 for vertical skew. However, particularly in the large UK Biobank dataset, the significance levels were unambiguous despite the subtle effects (*p* value as low as 5.00e-20 for handedness with vertical skew).

As regards the horizontal component, previous results with regard to handedness have been mixed, with some studies finding an association of handedness with brain torque [2, 31–33], and others not [36–39]. All three of our datasets showed the same direction of effect for the association between handedness and horizontal skew, although limited statistical power to detect this effect in the BIL&GIN dataset (*N* = 453) is likely to explain that the association was not significant in this specific dataset. For the vertical skew, the association with handedness was again consistent in direction across all three datasets, and also significant in all three datasets. The association of handedness with the vertical component of global brain asymmetry is a novel finding, as far as we are aware. Given the small effect sizes in our study, it is clear that inferring the handedness of individuals from their global brain asymmetry is unlikely to be possible. Anatomists and anthropologists have long noted a potential link between left-handedness and brain asymmetry, which was initially considered to involve localized thinning and protrusions of the skull, such that attempts have even been made to use skull endocasts to infer the evolution of handedness in hominins [2, 34, 35, 40]. For example, based on their asymmetrical shapes, a skull from Gibraltar and the ‘Peking man’ were suggested to be from right-handed individuals, while a skull from London was suggested to have been from a left-handed individual (for a review, see [40]). Unfortunately, relationships between brain structural and functional asymmetries are clearly far from absolute, as evidenced by the present study, as well as others (e.g., [37–39]).

We found no significant genetic correlations between either aspect of global brain asymmetry and handedness in the UK Biobank or HCP datasets. This suggests that genetic factors influencing global brain asymmetry are largely dissociable from those affecting handedness, and therefore that environmental factors, such as early life experiences, may play a more predominant role in causing the associations of handedness with global brain asymmetry measures.

### Development of global brain asymmetry

Thus far we know little about the developmental mechanisms which lead to brain asymmetry. Best (1986) proposed a 3D lateralized neuro-embryologic growth gradient, including a “rearward and dorsal” twist of the left hemisphere and a “forward and ventral” twist of the right hemisphere. However, the dorsal-ventral twist was based only on a preliminary observation at the time [92], and our observations of adults in the present study showed an opposite direction, i.e. a ventral twist of the left hemisphere, and a dorsal twist of the right. However, the typical adult brain asymmetries are the endpoint of a dynamic developmental process that also plays out through childhood and adolescence, and may involve some reconfiguration [20]. Possible mechanisms may include inter-hemispheric differences in neural pruning [21], axon tension [93], and/or ventricular cerebrospinal fluid (CSF) volume [22] during neural development.

Establishing the genetic contributions to brain asymmetry would help elucidate the developmental origins of this trait, as well as potentially its evolution, and the neural basis of functional lateralization. In the present study, we found heritabilities of 4-13% for the two global brain asymmetry measures in the UK biobank and HCP datasets. This is lower than the 10%-30% heritabilities reported in a previous study of vervet monkeys using a similar registration-based approach [12], which might suggest a species difference in the degree of genetic control of global brain asymmetry [8].

Our GWAS analyses revealed no loci that surpassed the standard genome-wide significance threshold *p*<5e-08, for either horizontal or vertical skew. Other recent GWAS studies [49–51], also using UK Biobank data, were able to identify significant loci affecting various regional asymmetries (cortical regional surface area and thickness asymmetries, and subcortical volume asymmetries), which had heritabilities in the same range as the skew measures in the present study. In addition, both skews showed high measurement reliability in twice-scanned individuals. The lack of significant GWAS results with the skew measures was therefore likely due to high polygenicity, as was indicated by causal mixture model analysis in the UK Biobank data. This is also consistent with the associations of skew measures with various different regional grey and white matter metrics, many of which can have distinct genetic contributions [94, 95], such that the genetic architecture of global asymmetry measures is likely to be particularly complex. Even larger GWAS analyses will be required to pinpoint significant SNPs associated with brain skews.

Our gene-based analysis, in which SNP-level association statistics were combined at the gene-level for a single test per gene, did not identify individual genes at a statistically significant level after adjustment for multiple testing. The most significant gene *PLEC* encodes plectin, which is a cytoskeletal protein linking the three main components of the cytoskeleton: actin microfilaments, microtubules and intermediate filaments [96]. Cytoskeletal-related genes have been implicated by other, recent genetic studies of regional structural brain asymmetry and handedness, as well as functional hemispheric language dominance [50, 52–54, 97]. Cytoskeletal-mediated mechanisms for left-right asymmetry development have also been described in invertebrates and frogs [55–58, 97]. We have therefore previously proposed the existence of a human brain-intrinsic mechanism of left-right axis determination, involving cytoskeletal influences on cellular chirality, which may be developmentally distinct from left-right laterality of the visceral organs [97].

In addition, gene-set enrichment analysis of the GWAS results for vertical skew tentatively implicated genes involved in neuron projection guidance. This may relate, for example, to the establishment of interhemispheric connections via the corpus callosum. Consistent with this hypothesis, a recent GWAS analysis of publicly released, imaging-derived phenotypes in the UK Biobank [98] (https://open.win.ox.ac.uk/ukbiobank/big40/pheweb/, database queried on August 1, 2020) found that corpus callosum measures (e.g., fractional anisotropy of the genu of the corpus callosum, and volume of the anterior part of middle corpus callosum) showed the most significant associations annotated to *PLEC*.

As the heritabilities of both global asymmetry measures were low in our analyses, then non-genetic factors also seem likely to influence them. Early life factors such as birth weight, being part of a multiple birth, and breastfeeding, have been shown to correlate with handedness [29]. In the present study, while we did not find any significant associations between global brain asymmetries and early life factors after correction for multiple testing, two nominally significant associations were observed with vertical asymmetry: breastfeeding and birth weight. In addition, country of birth was significant even after correction for multiple testing in phenome-wide association analysis. There may therefore be aspects of prenatal and perinatal behaviour, or care, which differ systematically between the countries of the United Kingdom, and can affect brain shape development. Population-genetic differences may also be involved, as may environmental influences later in life.

### Other notable phenotypic and genetic correlations with brain asymmetry skews

As abnormal brain asymmetry patterns have been reported in a variety of cognitive and neuropsychiatric disorders, including dyslexia [15], schizophrenia [80], attention-deficit/hyperactivity disorder [20], autism [99], obsessive-compulsive disorder [100], the two measures of global asymmetry used in the present study might usefully be analyzed in future studies of these disorders. In the UK Biobank, there were significant phenotypic correlations of both horizontal and vertical brain skew with a depression-related variable, ‘*Recent changes in speed/amount of moving or speaking*’. This may be consistent with previous reports of altered occipital bending in major depression [21, 22]. Within the HCP dataset, there was a positive correlation between horizontal skew and oral reading recognition ability. Language-related cognition is well known to make use of lateralized functional networks [101, 102]. In addition, we found evidence for genetic correlation between horizontal brain skew and ASD, which may be consistent with the subtle but widespread alterations of regional cortical thickness asymmetry in ASD reported in a recent, large-scale study [79]. The automated measurement of global brain asymmetry that we have employed here will be feasible for large-scale meta-analysis-based studies of brain disorders, such as those carried out within consortia such as ENIGMA (http://enigma.ini.usc.edu/) [103].

In addition, we found horizontal and vertical brain skew measures to show significant phenotypic correlations with body mass index in the UK Biobank. This is consistent with a recent observation that body mass index correlated with regional thickness of frontal and occipital regions, in opposite directions in the two hemispheres, in a study of 895 healthy adults [104]. Further studies will be need to disentangle any potential cause-effect relations and underlying mechanisms linking these traits. For the pheWAS analysis with brain skew measures, we adjusted for various covariate effects: sex, age, nonlinear age, the first ten principal components that capture genome-wide population structure in the genotype data, and technical variables related to imaging (imaging assessment center, scanner position parameters, and signal/contrast-to-noise ratio in T1). Nonetheless, other confounding effects may have been relevant for the associations with some phenotypes. PheWAS is a screening approach, applied under a single model with fixed covariates, across thousands of phenotypes. In principle, any one of the thousands of phenotypes could be a relevant confounder for any of the others when assessing its relation with brain skew. There may also be unmeasured, underlying effects that cause some combinations of traits to be associated. For these reasons, we make no claims about cause-effect relations based on the PheWAS results.

### Conclusion

In sum, the present study used automated, registration-based measurement of the horizontal and vertical components of global brain asymmetry, both of which showed high test-retest repeatability. With the largest-ever analyses, we revealed two average asymmetry patterns at the population level: one global asymmetry along the anterior-posterior axis, and one along the dorsal-ventral axis. Furthermore, we clarified the relationships between global brain asymmetries and handedness, linking brain structural asymmetries to one of the most clearly evident functional lateralizations, although the effect sizes were small. The two asymmetrical skew measures also showed associations with diverse metrics of regional grey matter and white matter, as well as various phenotypic variables related to cognitive functions, sociodemographic and physical factors, and mental health. Genetic analyses indicated low heritability and high polygenicity of the brain skew measures, and a potential genetic overlap with ASD. Together, our results provide evidence for the functional significance of global brain asymmetry, and indicate that genetic variation plays a role - although not a determining one - in its variation in the population.

## Ethics statement

This study utilized de-identified data from the baseline and imaging assessments of the UK Biobank, a prospective cohort of 500,000 individuals (age 40-69 years) recruited across Great Britain during 2006-2010. The protocol and consent were approved by the UK Biobank’s Research Ethics Committee. Data from the Human Connectome Project was approved by the Institutional Review Boards associated with that project. The BIL&GIN study was approved by the local ethics committee (CCPRB Basse-Normandie).

## Data Availability

For use of UK Biobank data, application must be made via http://www.ukbiobank.ac.uk/register-apply/. The Human Connectome Project data are available via https://www.humanconnectome.org/. The BIL&GIN data sharing is based on a collaborative model: http://www.gin.cnrs.fr/BIL&GIN.

## Code Availability

All analyses were carried out as described in the methods. Software versions and relevant parameters are included in the corresponding methods sections. Scripts are available from the authors upon request.

## Acknowledgements

This research was funded by the Max Planck Society (Germany) and grants from the Netherlands Organization for Scientific Research (NWO) (054-15-101) and the French National Research Agency (ANR, grant No. 15-HBPR-0001-03), as part of the FLAG-ERA consortium project ‘MULTI-LATERAL’, a Partner Project to the European Union’s Flagship Human Brain Project. We thank Nathalie Tzourio-Mazoyer for sharing her extensive knowledge of brain laterality. This research was conducted using the UK Biobank resource under Application Number 16066, with Clyde Francks as the principal applicant. Our study made use of imaging-derived phenotypes generated by an image-processing pipeline developed and run on behalf of UK Biobank. We thank the UK Biobank and the Human Connectome Project for data sharing.

## Author contributions

X.Z.K. and C.F. designed the research. X.Z.K, M.P., A.P., and A.C.C performed data analysis; X.Z.K. and C.F. drafted the paper. All authors provided advice on the study and feedback on the paper.

## Notes

### Competing Interest Statement

The authors have declared no competing interest.

### Summary of Updates

An issue in the software MAGMA has been fixed recently. In this revision, we have updated all relevant results using the latest version MAGMA.

