## Supplemental S for "Large-scale Phenomic and Genomic Analysis of Brain Asymmetrical Skew"

**Contents:**

**Figure S1-S8**

**Tables S1-S5**

**Dataset S1-S2 (separate files)**

**Fig. S1. Examples of two subjects with different scores for horizontal skew.** The red arrows indicate the skewing of each brain during image registration.

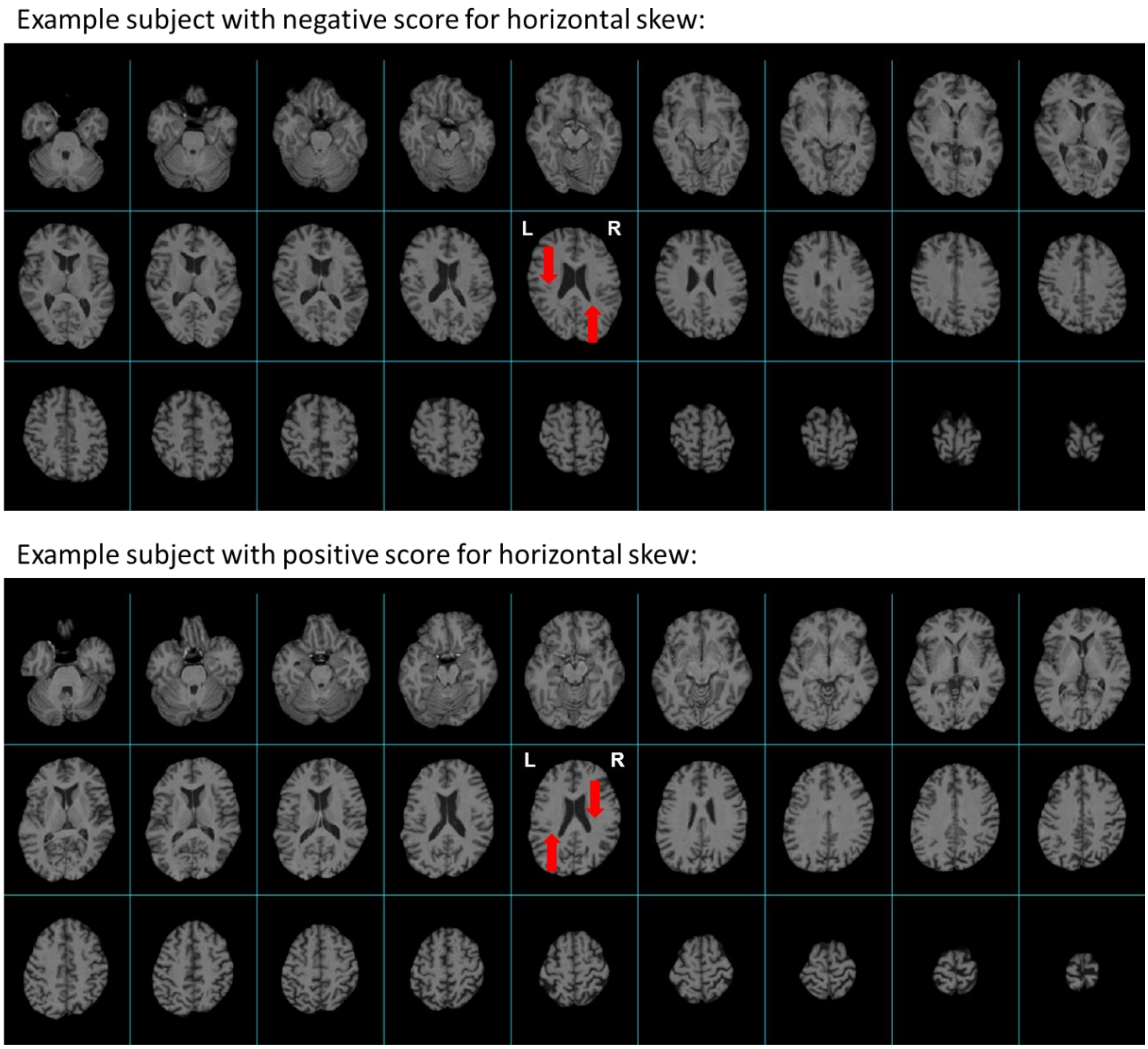

**Fig. S2. Examples of two subjects with different scores for vertical skew.** The red arrows indicate the skewing of each brain during image registration.

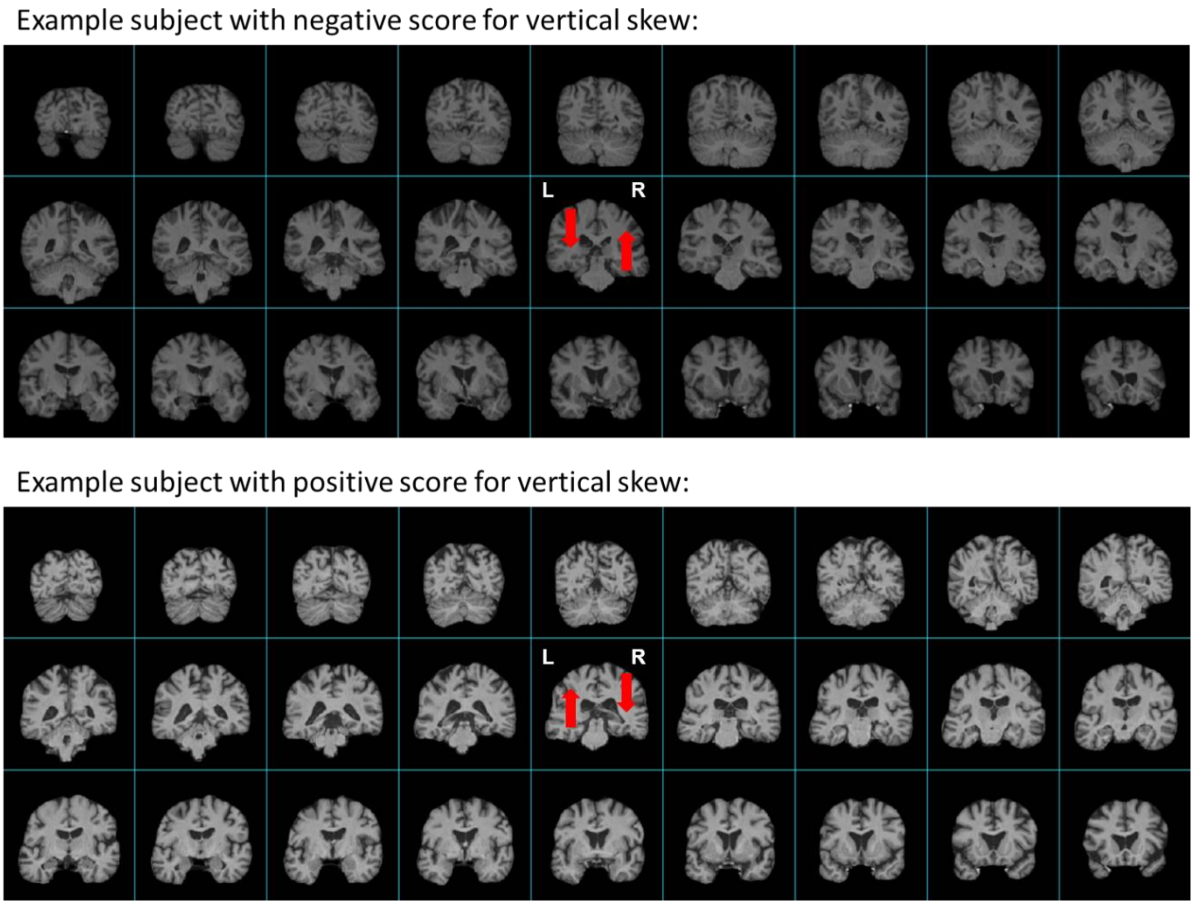

**Fig. S3. QQ-plots for pheWAS of horizontal skew and vertical skew.** Red stars indicate that P values are smaller than 5e-324 (the smallest possible on a typical R platform).

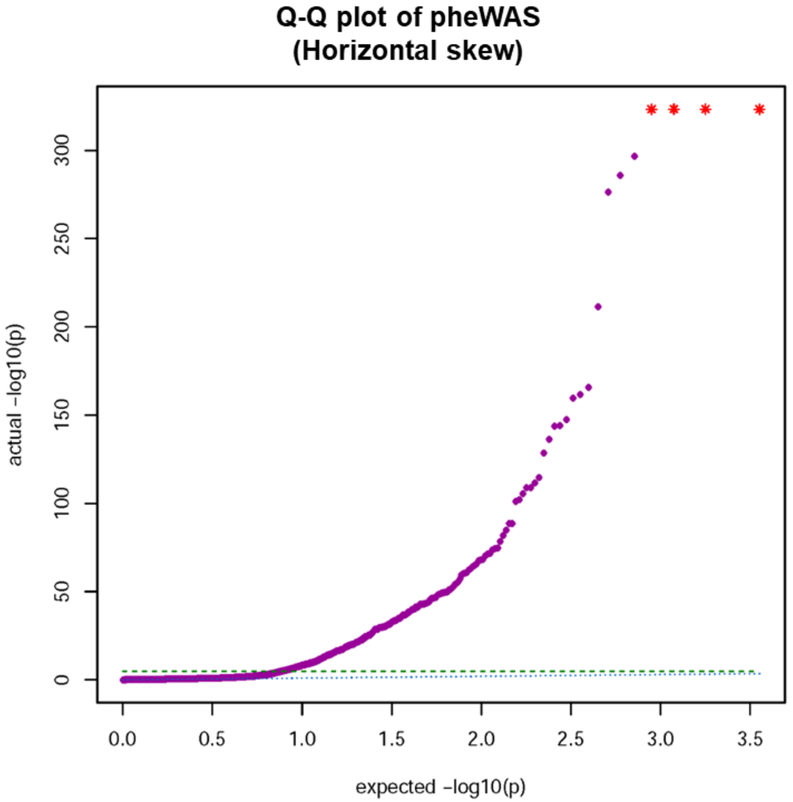

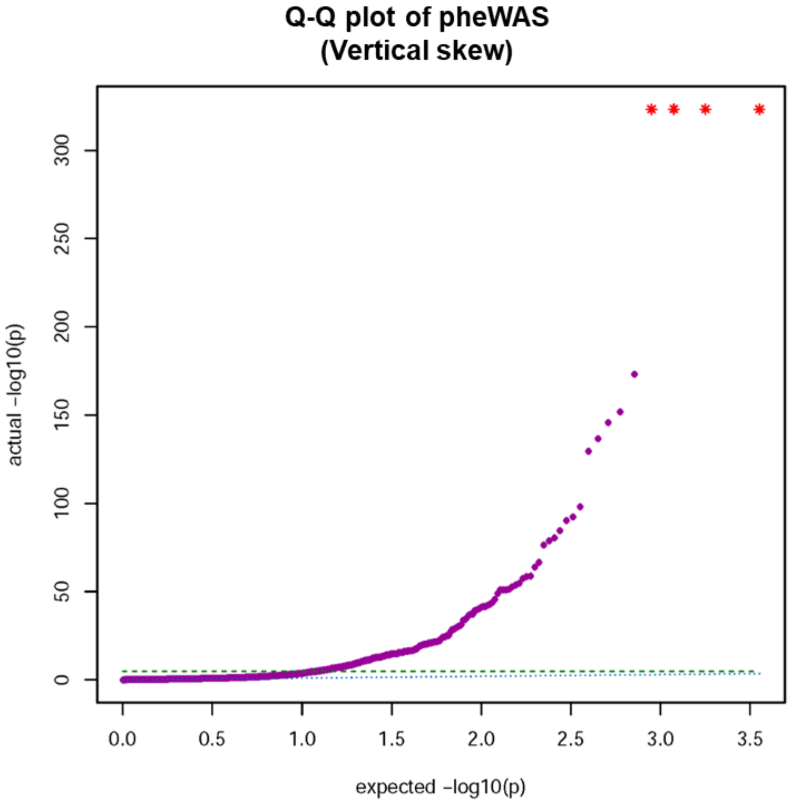

**Fig. S4. Manhattan Plots for GWAS analyses.** (A) Horizontal skew. (B) Vertical skew. No loci surpassed the genome-wide significance threshold (5e-08).

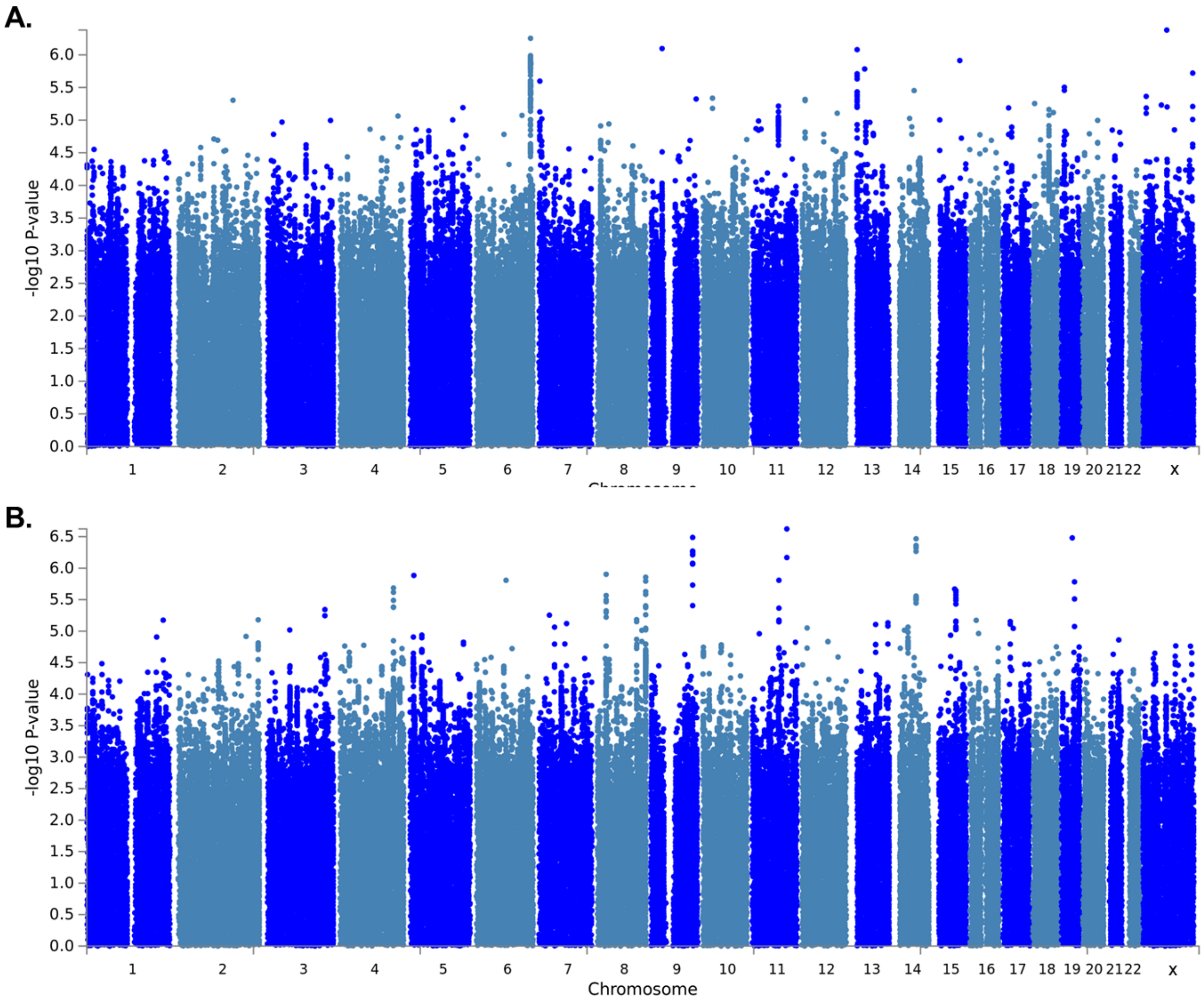

**Fig. S5. Association results of the asymmetrical skews with regional grey matter volumes. Red indicates a positive association; blue indicates a negative association.** (A) Horizontal skew. (B) Vertical skew.

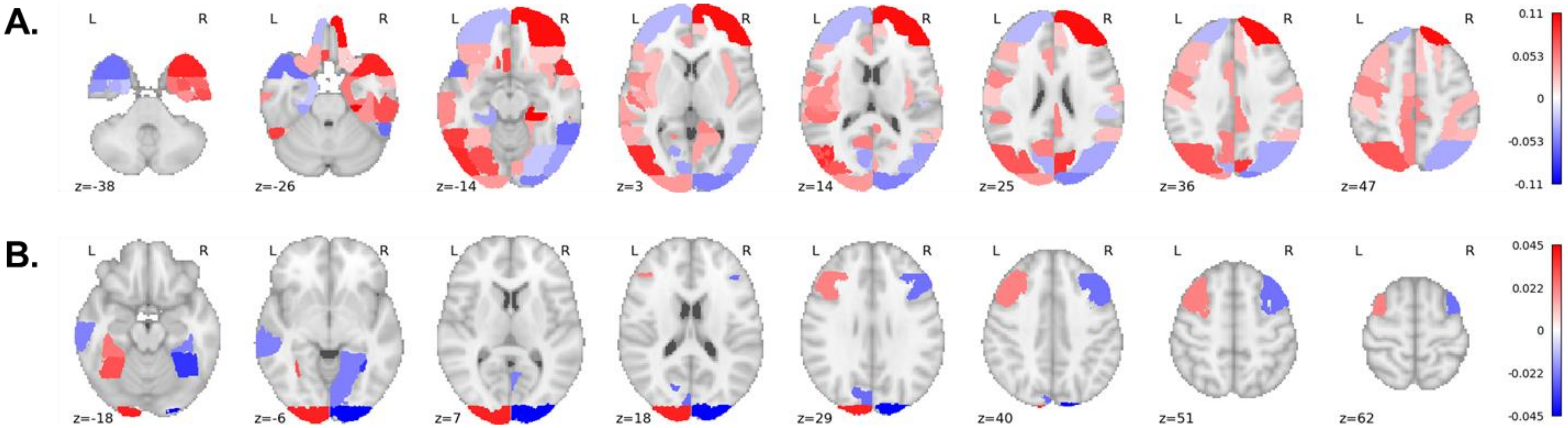

**Fig. S6-1. Association results of the horizontal skew with various white matter metrics.** Red indicates a positive association; blue indicates a negative association. FA: fractional anisotropy, MD: mean diffusivity, L1/ L2/L3: the three eigenvalues of diffusion, MO: mode of anisotropy, OD: orientation dispersion, ICVF: intra-axonal volume fraction, ISOVF: isotropic volume fraction.

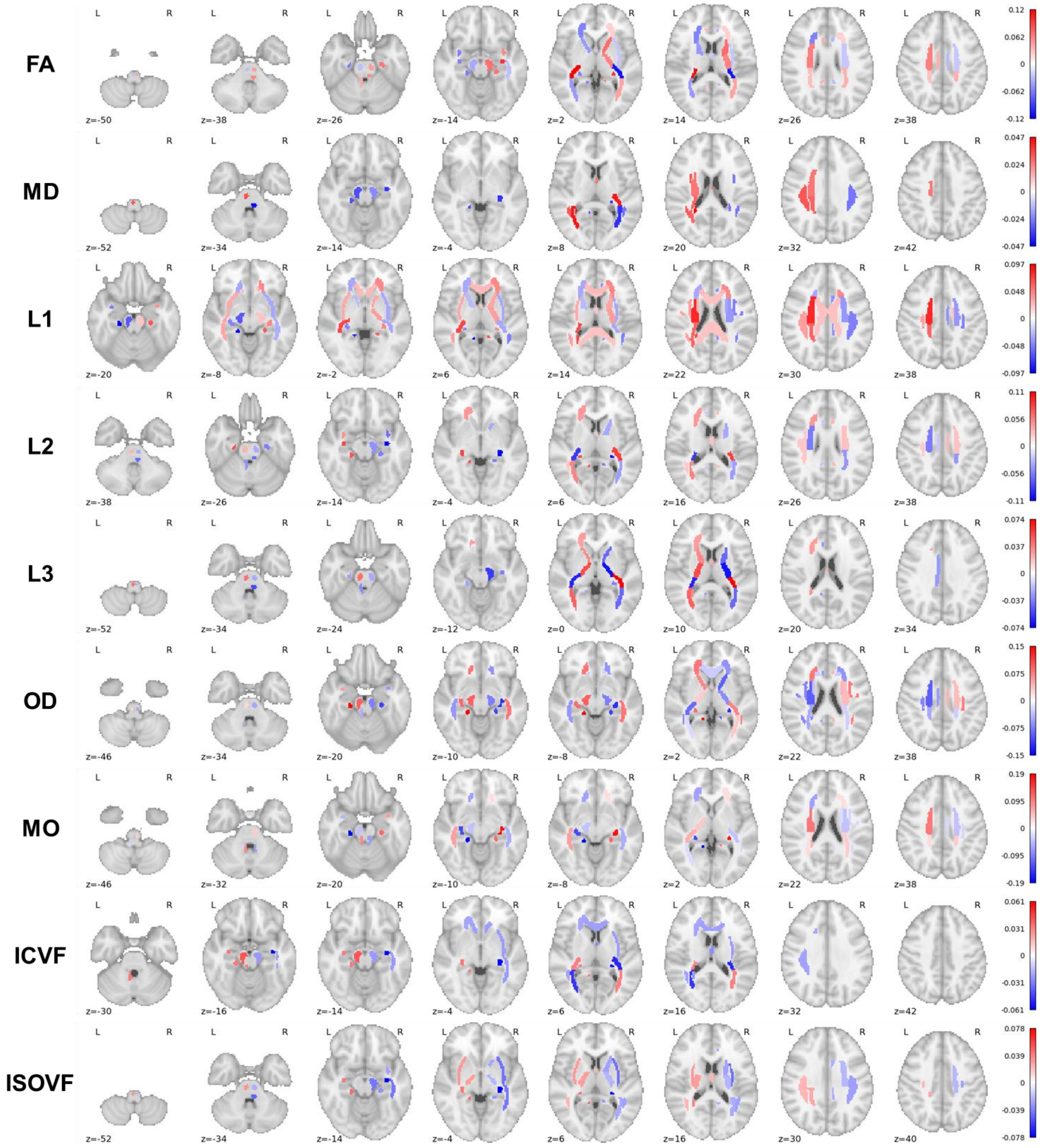

**Fig. S6-2. Association results of the horizontal skew with various white matter metrics (FA weighted).** Red indicates a positive association; blue indicates a negative association. FA: fractional anisotropy, MD: mean diffusivity, L1/ L2/L3: the three eigenvalues of diffusion, MO: mode of anisotropy, OD: orientation dispersion, ICVF: intra-axonal volume fraction, ISOVF: isotropic volume fraction.

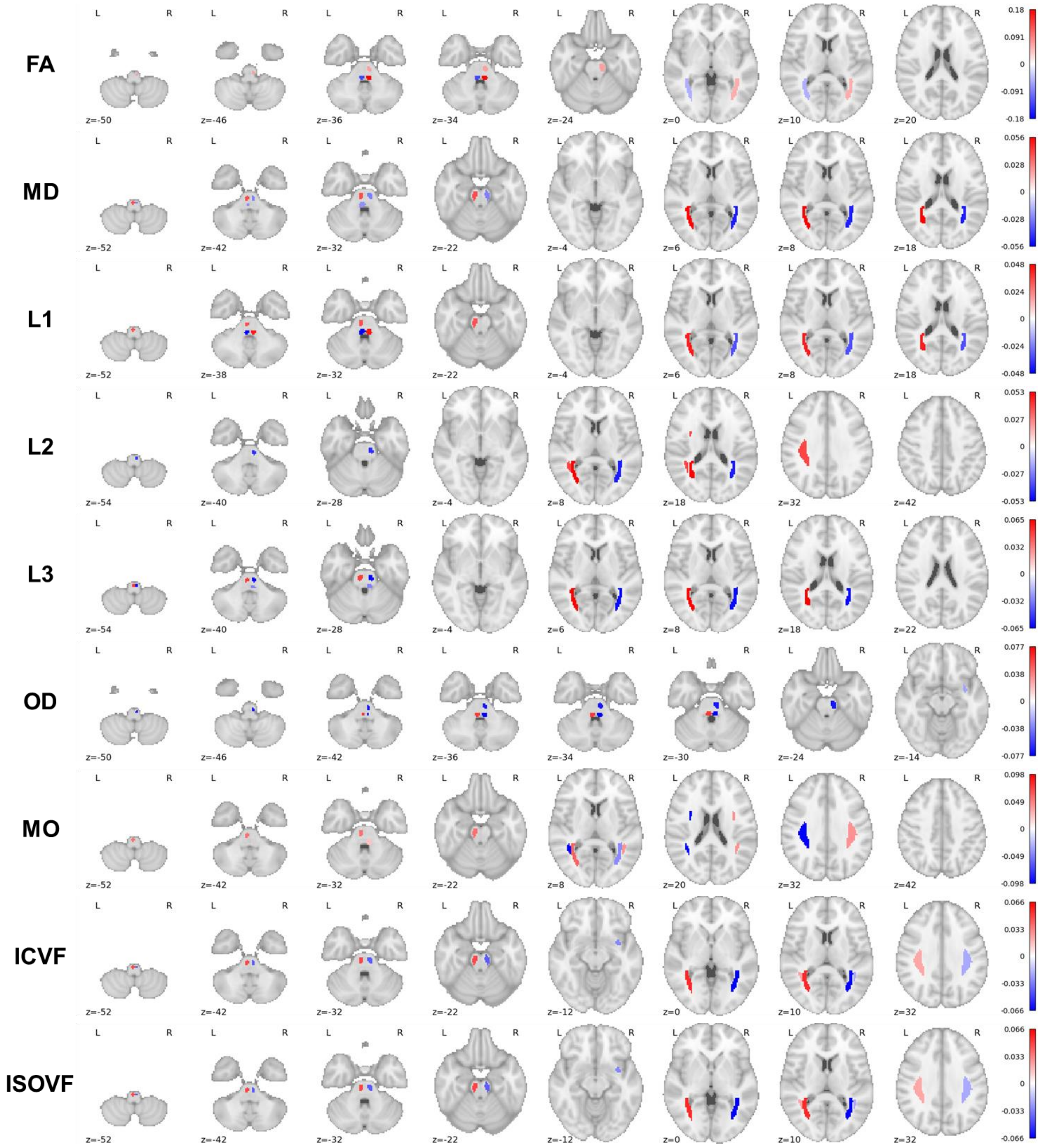

**Fig. S7-1. Association results of the horizontal skew with various white matter metrics.** Red indicates a positive association; blue indicates a negative association. FA: fractional anisotropy, MD: mean diffusivity, L1/ L2/L3: the three eigenvalues of diffusion, MO: mode of anisotropy, OD: orientation dispersion, ICVF: intra-axonal volume fraction, ISOVF: isotropic volume fraction.

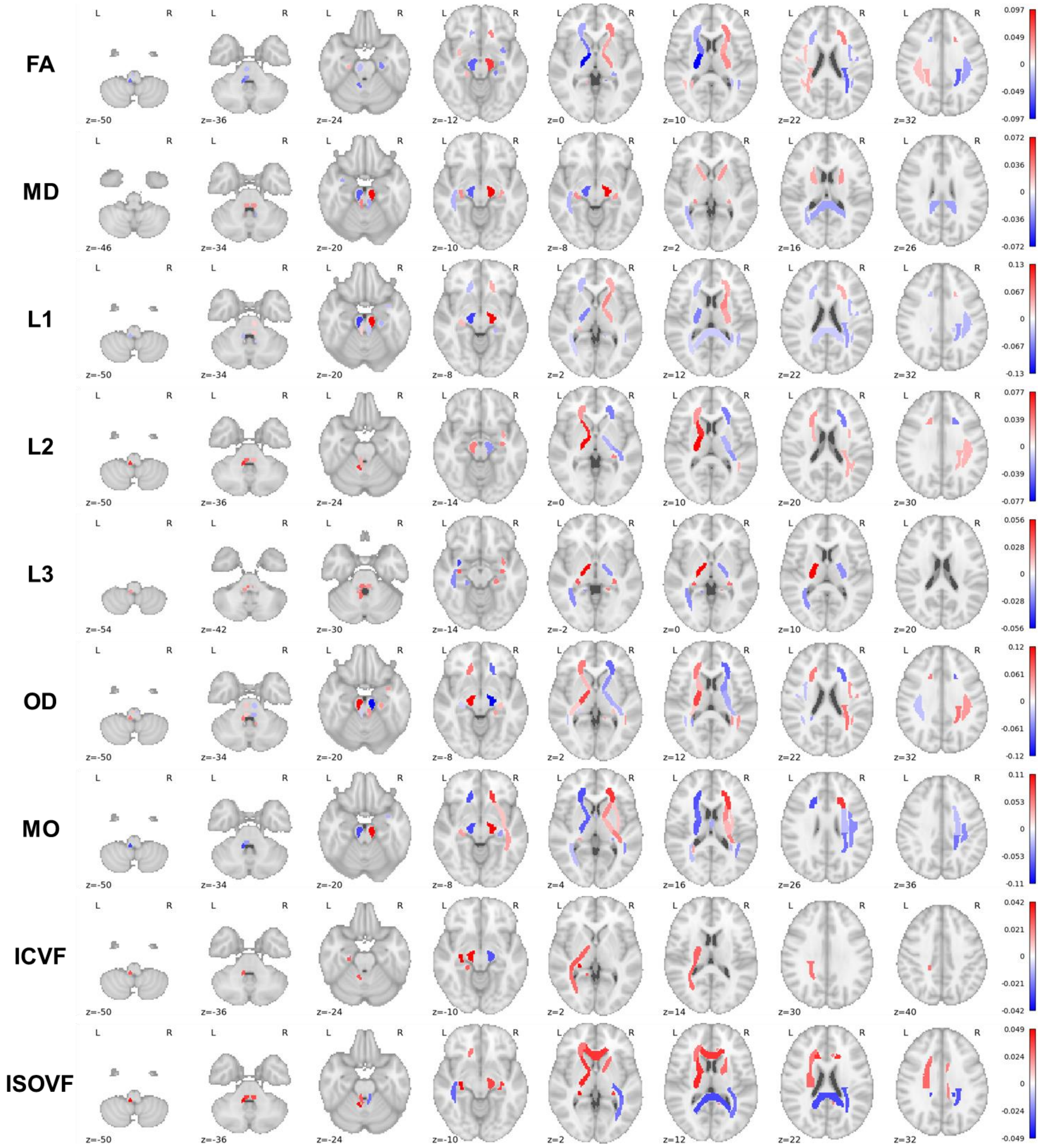

**Fig. S7-2. Association results of the horizontal skew with various white matter metrics (FA weighted).** Red indicates a positive association; blue indicates a negative association. FA: fractional anisotropy, MD: mean diffusivity, L1/ L2/L3: the three eigenvalues of diffusion, MO: mode of anisotropy, OD: orientation dispersion, ICVF: intra-axonal volume fraction, ISOVF: isotropic volume fraction.

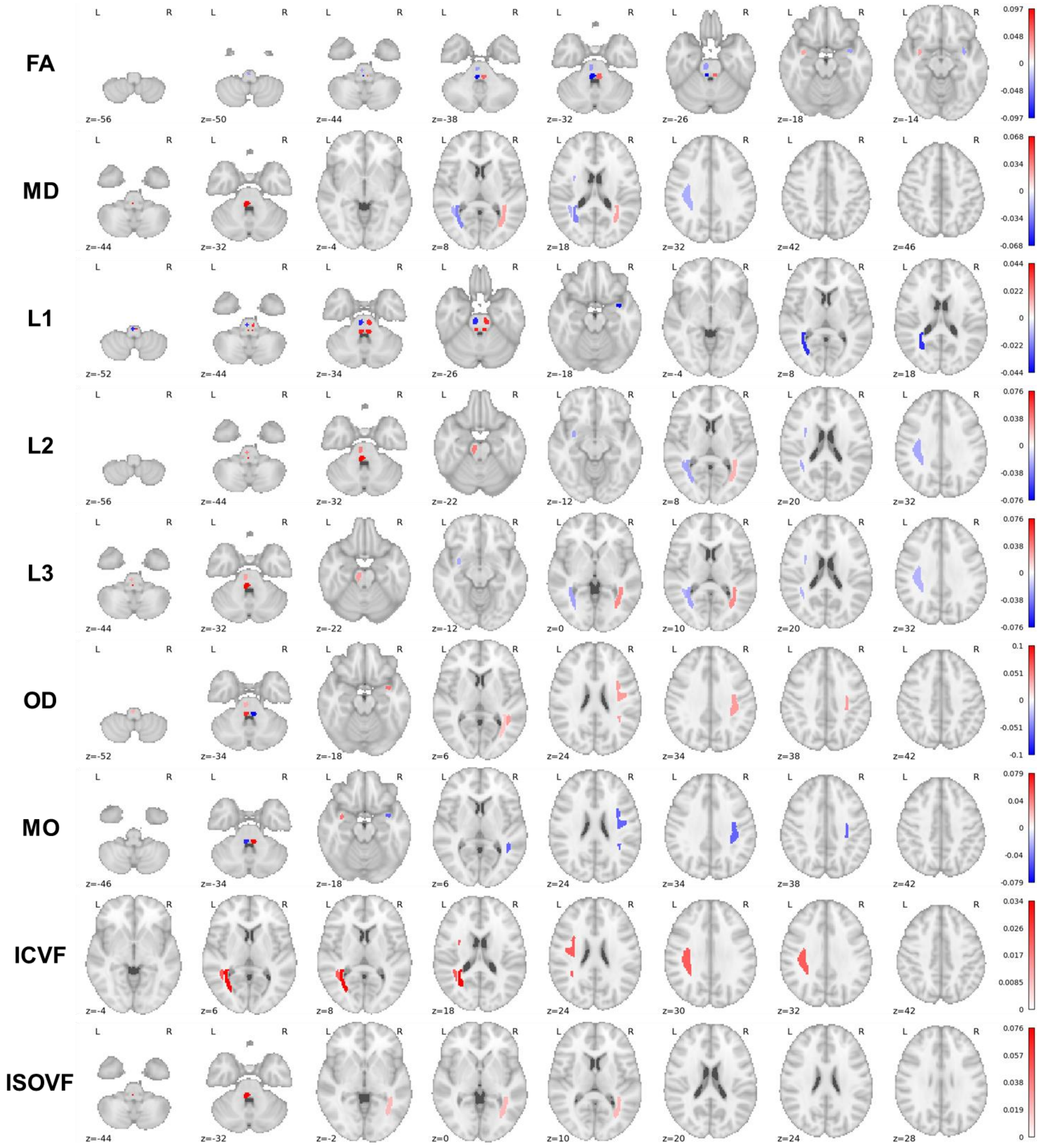

**Table S1. Correlations between the skew measures and brain-size-related scaling factors in the three datasets.** Correlations in bold indicate correlations that could survive correlation for multiple testing ($p < 0.002$, i.e., 0.05/24).

| | *Dataset* | *ScalesAvg* | | *ScalesX* | | *ScalesY* | | *ScalesZ* | |
|---|---|---|---|---|---|---|---|---|---|
| | | *r* | *p* | *r* | *p* | *r* | *p* | *r* | *p* |
| *Horizontal skew* | UKB | **-0.06924** | **2.25E-43** | **-0.06002** | **5.35E-33** | **-0.04681** | **1.06E-20** | **-0.05587** | **8.34E-29** |
| | HCP | -0.0829 | 0.005652 | **-0.1024** | **0.000623** | -0.05295 | 0.077454 | -0.05529 | 0.065195 |
| | BIL | -0.07254 | 0.123155 | -0.03303 | 0.483111 | -0.09676 | 0.039538 | -0.04369 | 0.353551 |
| *Vertical skew* | UKB | **-0.02216** | **1.01E-05** | -0.01125 | 0.025077 | -0.01534 | 0.002252 | **-0.02422** | **1.40E-06** |
| | HCP | -0.08127 | 0.006671 | -0.04652 | 0.120863 | **-0.11682** | **9.37E-05** | -0.04419 | 0.140692 |
| | BIL | -0.04986 | 0.289666 | -0.05653 | 0.22982 | -0.0391 | 0.406452 | -0.02471 | 0.599831 |

**Table S2. SNPs suggestively associated with brain skews (association *p* values less than 5e-07 in GWAS).**

| | *rsid* | *chr* | *pos* | *Effect Allele* | *Other Allele* | *MAF* | *info* | *beta* | *se* | *t* | *P* | *Nearest gene* |
|---|---|---|---|---|---|---|---|---|---|---|---|---|
| *Horizontal* | rs147003578 | X | 73839775 | A | C | 0.026426 | 0.94317 | 0.0026167 | 0.00051705 | 5.0608 | 4.17E-07 | *RLIM* |
| *Vertical* | rs117457382 | 11 | 103798827 | A | G | 0.02179 | 0.99837 | -0.0020824 | 0.00040303 | -5.167 | 2.38E-07 | *PDGFD* |
| | rs10136315 | 14 | 69063564 | A | G | 0.40092 | 0.99985 | -0.00060738 | 0.00011912 | -5.099 | 3.42E-07 | *RAD51B* |
| | rs138596065 | 19 | 33010000 | G | GC | 0.081832 | 0.98091 | -0.0011546 | 0.00022616 | -5.1051 | 3.30E-07 | *DPY19L3* |
| | rs113249637 | 9 | 124908299 | G | A | 0.05362 | 0.99882 | 0.001336 | 0.00026154 | 5.1084 | 3.25E-07 | *NDUFA8* |

**Table S3. Top gene ontology terms in the gene-set analyses for each of the asymmetrical skews (*p* less than 0.001).** Gene ontology terms **in bold** survive correction of multiple testing for a given skew measure (*p* = 7.60e-06, i.e., 0.05/6576).

| | *FULL_NAME* | *NGENES* | *BETA* | *BETA_STD* | *SE* | *P* |
|---|---|---|---|---|---|---|
| *Horizontal Skew* | GO_bp:go_interaction_with_symbiont | 78 | 0.32279 | 0.020044 | 0.092889 | 0.000256 |
| | GO_bp:go_sterol_metabolic_process | 96 | 0.29181 | 0.020094 | 0.085193 | 0.000308 |
| | GO_mf:go_e_box_binding | 51 | 0.39827 | 0.020011 | 0.11677 | 0.000325 |
| | GO_bp:go_regulation_of_neurotrophin_trk_receptor_signaling_pathway | 13 | 0.86336 | 0.021922 | 0.27278 | 0.000777 |
| *Vertical Skew* | **GO_bp:go_neuron_projection_guidance** | **272** | **0.21418** | **0.024715** | **0.04949** | **7.59E-06** |
| | GO_bp:go_cell_morphogenesis_involved_in_neuron_differentiation | 575 | 0.12548 | 0.020892 | 0.034548 | 1.41E-04 |
| | GO_bp:go_cell_morphogenesis_involved_in_differentiation | 726 | 0.10924 | 0.020359 | 0.030805 | 0.000196 |
| | GO_bp:go_glutathione_metabolic_process | 40 | 0.56254 | 0.025039 | 0.15948 | 0.000211 |
| | GO_bp:go_cell_part_morphogenesis | 673 | 0.10674 | 0.019178 | 0.031662 | 0.000375 |
| | GO_bp:go_axon_development | 504 | 0.12316 | 0.019233 | 0.03659 | 0.000382 |
| | GO_bp:go_negative_regulation_of_vascular_endothelial_growth_factor_receptor_signaling_pathway | 13 | 0.73254 | 0.0186 | 0.22497 | 0.000566 |

**Table S4. Top genetic correlations of the asymmetrical skews with traits in the UK Biobank.** Uncorrected p < 0.01.

| *Skew* | *Trait Name* | *Trait Field ID* | *Trait Category* | *beta* | *se* | *z* | *p* |
|---|---|---|---|---|---|---|---|
| *Horizontal Skew* | Volume of grey matter in I-IV Cerebellum (left) | f.25893.2.0 | T1 structural brain MRI | 0.350074 | 0.138952 | 2.519388 | 0.005878 |
| *Horizontal Skew* | Volume of grey matter in X Cerebellum (right) | f.25920.2.0 | T1 structural brain MRI | 0.431539 | 0.168821 | 2.556193 | 0.005291 |
| *Horizontal Skew* | Volume of grey matter in Crus II Cerebellum (left) | f.25903.2.0 | T1 structural brain MRI | 0.327265 | 0.132636 | 2.467392 | 0.006805 |
| *Horizontal Skew* | Mean MO in uncinate fasciculus on FA skeleton (left) | f.25197.2.0 | Diffusion brain MRI | -0.38483 | 0.160576 | -2.39657 | 0.008275 |
| *Horizontal Skew* | Mean L2 in uncinate fasciculus on FA skeleton (left) | f.25293.2.0 | Diffusion brain MRI | 0.363293 | 0.148064 | 2.453621 | 0.007071 |
| *Horizontal Skew* | Volume of grey matter in Heschl's Gyrus (includes H1 and H2) (left) | f.25870.2.0 | T1 structural brain MRI | 0.364242 | 0.154645 | 2.355343 | 0.009253 |
| *Vertical Skew* | Mean OD in cerebral peduncle on FA skeleton (left) | f.25407.2.0 | Diffusion brain MRI | 0.207555 | 0.083014 | 2.500241 | 0.006205 |
| *Vertical Skew* | Place of birth in UK - east co-ordinate | f.130.0.0 | Early life factors | -0.22382 | 0.092396 | -2.42243 | 0.007709 |

**Dataset S1. pheWAS results for horizontal skew after multiple testing correction (*p* less than 1.40e-05, i.e., 0.05/3562).**

See the attached table.

**Dataset S2. pheWAS results for vertical skew after multiple testing correction (*p* less than 1.40e-05, i.e., 0.05/3562).**

See the attached table.
